# TRACKING HEMATOPOIETIC STEM CELL EVOLUTION IN A WISKOTT-ALDRICH CLINICAL TRIAL

**DOI:** 10.1101/2022.05.30.494052

**Authors:** Danilo Pellin, Luca Biasco, Serena Scala, Clelia Di Serio, Ernst C. Wit

## Abstract

Hematopoietic Stem Cells (HSC) are the cells that give rise to all other blood cells and, as such, they are crucial in the healthy development of individuals. Wiskott-Aldrich Syndrome (WAS) is a severe disorder affecting the regulation of hematopoietic cells and is caused by mutations in the WASP gene. We consider data from a revolutionary gene therapy clinical trial, where HSC harvested from 3 WAS patients’ bone marrow have been edited and corrected using viral vectors. Upon re-infusion into the patient, the HSC multiply and differentiate into other cell types. The aim is to unravel the cell multiplication and cell differentiation process, which has until now remained elusive.

This paper models the replenishment of blood lineages resulting from corrected HSC via a multivariate, density-dependent Markov process and develops an inferential procedure to estimate the dynamic parameters given a set of temporally sparsely observed trajectories. Starting from the master equation, we derive a system of non-linear differential equations for the evolution of the first- and second-order moments over time. We use these moment equations in a generalized method-of-moments framework to perform inference. The performance of our proposal has been evaluated by considering different sampling scenarios and measurement errors of various strengths using a simulation study. We also compared it to another state-of-the-art approach and found that our method is statistically more efficient.

By applying our method to the Wiskott-Aldrich Syndrome gene therapy data we found strong evidence for a myeloid-based developmental pathway of hematopoietic cells where fates of lymphoid and myeloid cells remain coupled even after the loss of erythroid potential.

All code used in this manuscript can be found in the online Supplement, and the latest version of the code is available at github. com/dp3ll1n/SLCDP_v1.0.

## 1. Introduction

Although mammalian organisms have more than a hundred different cell types, many tissues are sustained by relatively few varieties of multipotent stem and progenitor cells (Weissman, 2000; Blanpain, Horsley and Fuchs, 2007; Snippert and Clevers, 2011). Given their importance, a comprehensive understanding of stem cells is crucial for advancing the development of regenerative medicine. HSC represent a particular pool of cells that resides mainly in the bone marrow and has the unique capability of self-renewal. Through a process of progressive specialization called hematopoiesis, HSC can give rise and replenish all blood lineages in a human being, lifelong. HSC are among the most clinically relevant cell population and are used to treat many hematological malignancies and bone marrow disorders. Despite being the focus of decades of research and clinical efforts, many questions about HSC biology are still open and debated. For example, it is well-established that a progressive loss of multi-lineage potential occurs when descending the hematopoietic cell differentiation hierarchy from HSC to committed cell types and then, finally, mature blood cells. However, it is still unclear at what stage of the differentiation process the separation between the three main cell lineage groups, lymphoid, myeloid, and erythroid, happens. Other essential aspects about the metabolism of human blood cells, such as how duplication, death, and differentiation rates are orchestrated along the blood phylogeny to maintain the hematopoietic system stable, are still unknown.

Gene therapy consists of delivering DNA or RNA fragments into cells of patients as a drug to treat a disease. It has been mainly applied to inherited monogenetic disease where deleterious mutations occurring in a specific known gene lead to the synthesis of a dysfunctional protein causing the symptoms. Under this setting, gene therapy offers a real opportunity and can be used to provide cells with a correct copy of the gene, thereby producing a functional version of the protein. The treatment effect is tied to the presence and activity of the therapeutic gene in specific cells or tissue, hence for the long-term treatment of hematological disorders, HSC represent the ideal target for gene therapy clinical trials (Naldini, 2011; Biffi et al., 2013; Aiuti et al., 2013).

This paper will focus on a gene therapy clinical trial for Wiskott-Aldrich Syndrome (WAS), an inherited immunodeficiency caused by mutations in the gene encoding for WAS protein. The study was performed by the authors of this paper and described in clinical detail in Biasco et al. (2016). Briefly, HSC sorted from patient’s bone marrow samples according to their immunophenotyping characteristic — enrichment analysis for known protein on a cell’s cellular membrane, such as CD34 molecules specifically for HSC isolation — are distinctly labeled through the random incorporation of the WASP gene into their genome, using a lentiviral vector. Importantly, all progeny deriving from a marked HSC, through both duplication and differentiation, will carry the corrected copy of the gene and the identical unique markings defined by the original viral insertion site (IS). This procedure allows not only to obtain a long-term and widespread expression of the WAS protein among all blood lineages but also to perform in-vivo clonal tracking, the longitudinal observation of multiple clones’ evolution. It is crucial to highlight that for ethical reasons, gene therapy is one of the few settings in which scientists can collect information about human, *in-vivo* hematopoiesis at clone level.

One of the first quantitative analysis of clonal tracking data was developed in the context of a non-human primate rhesus macaque study by Wu et al. (2014). Using clustering methods on the multi-lineage clonal output of barcoded HSC, authors demonstrated how the correlation among lineages changes during reconstitution, with uni-lineage short-term progenitors being supplanted over time by multi-lineage long-term clones. (Biasco et al., 2016; Pellin et al., 2019) model clones dynamics using local linear approximations. Assuming linearity offers several advantages from a computational perspective, but also implies that cell type counts must eventually either go to zero or infinity in the long term. This assumption is biologically unrealistic because the hematopoietic system evolves in a constrained environment with limited resources and space available. At the same time, the replenishment of blood cells lasts for the entire life span of a human being. To extrapolate insight from real data Biasco et al. (2016) and Pellin et al. (2019) relied on a first-order local linear approximation of the dynamics: this is efficient but not very accurate when the time between consecutive process measurements is large, as it is in the case of gene therapy clinical trials. Xu et al. (2019) re-analyzed the rhesus macaque data using a statistical framework that models hematopoiesis as a multi-type Markov branching process, similar to our set-up. In Xu et al. (2019), clone trajectories are considered realizations from a stochastic process defined using a set of fundamental cellular events with event-specific rates. The authors showed that it is possible to derive exact analytical formulation for the evolution of the moments through a set of ordinary differential equations (ODEs), given the cell differentiation tree configuration and assuming event rates to be linear in the cell counts. The estimation of the cell differentiation dynamic is performed by matching model-based correlation functions to empirical lineage temporal correlations. An alternative approach, similar to ours, could a be Bayesian implementation (Wilkinson, 2006; Golightly and Wilkinson, 2008), which can deal with temporal sampling an observational noise in a natural fashion. We expect that implementation of those methods would yield similar results to ours.

In section 2 we describe the clonal tracking data obtained from a gene therapy clinical trial for Wiskott-Aldrich syndrome, for which the statistical methodology in this paper has been developed. The stochastic cell differentiation process and its characteristics are presented in section 3. In section 4 a non-linear generalized least squares estimation procedure for the parameters in the stochastic process is developed, both from a methodological and computational point of view. section 5 is dedicated to simulation studies. In section 5.1 the performance of our proposal is compared for different sampling time intervals with a simpler polynomial generalized least squares estimation procedure. Section 5.2 and section 5.3 are focused respectively on the impact on inference performance of having (multiplicative) measurement errors on cell count observations and the effect of potential model misspecification. Section 5.4 compares our method to the correlation-based moment estimator by Xu et al. (2019). In section 6 we return to the WAS gene therapy clinical trial data and answer the main questions of this paper, namely, estimate the coefficients driving HSC differentiation and verify whether the WAS data support the classical dichotomy model or a recently proposed myeloid-based model of hematopoietic stem cell differentiation.

## 2. Hematopoietic stem cell gene therapy in Wiskott-Aldrich Syndrome patients

WAS syndrome is an X-linked primary immunodeficiency characterized by infections, micro-thrombocytopenia, eczema, autoimmunity, and lymphoid malignancies. The disorder is caused by mutations in the WAS gene, which encodes for WASP, a protein that regulates cytoskeleton conformation and is involved in proliferation, migration, and immunological synapsis formation. For patients without a matched donor, gene therapy based on the infusion of autologous gene-corrected HSC represent an alternative therapeutic strategy.

Three children with WAS, who did not have compatible allogeneic donors, were enrolled in phase I/II clinical trial. Autologous BM-derived CD34+ cells were collected, transduced with a lentiviral vector coding for human WASP under the control of a 1.6-kb reconstituted WAS gene promoter (LV-w1.6W) using an optimized protocol, and re-infused intravenously into the patients three days after collection. Patients are given chemotherapy treatment before receiving the engineered cell infusion to deplete the existing HSC compartment and to facilitate the engraftment of corrected cells. This conditioning procedure requires a fast replenishment of all blood lineages by corrected HSC upon infusion until a homeostasis condition is met. All three WAS patients showed robust and multi-lineage engraftment of gene-corrected cells in BM and PB up to the latest follow-up.

We collected IS from eight distinct Peripheral Blood (PB) and seven distinct Bone Marrow (BM) lineages at multiple time-points up to 36 months after infusion of transduced HSCs using a combination of linear-amplification-mediated (LAM)-PCR and next-generation sequencing (NGS) technologies (Biasco et al., 2011).

After initial clonal fluctuations, we observed stable and polyclonal reconstitution in all hematopoietic lineages starting from 1 year after the infusion of gene-corrected HSC. Importantly, no adverse event associated with insertional mutagenesis was detected, allowing us to exploit IS to assess hierarchical relationships among engineered blood cell types in humans.

A major distinction in three subgroups, named lymphoid, myeloid and erythroid branches, can be made within the hematopoietic cell types. The lymphoid branch, responsible for the adaptive immune system, can, in turn, be subdivided into T-cells (CD3 in BM and CD4, CD8, CD3 in PB), B-cell (CD19), and Natural Killer cells (NK-cells, CD56). Myeloid cell types are involved in such diverse roles as innate immunity, adaptive immunity, and blood clotting and are composed of monocytes (CD14), granulocytes (CD15), and megakaryocytes (CD61). Erythrocytes are the oxygen-carrying red blood cells (GLYCO).

Two different models of hematopoiesis are currently debated, shown in Figure 1. The classical dichotomy model assumes that HSC first generate a common myeloid-erythroid progenitor (CMEP) and a common lymphoid progenitor (CLP). The CLP then produces only T-cells or B-cells. The alternative myeloid-based model postulates that HSC first diverge into the CMEP and a common myeloid-lymphoid progenitor (CMLP), which generates T- and B-cell progenitors through a bipotential myeloid-T progenitor and a myeloid-B progenitor stage. The main difference is that according to the second, all erythroid, T- and B-lineage branches retain the potential to generate myeloid cells, even after the segregation of T- and B-cell lineages (Kawamoto, Wada and Katsura, 2010).

**Fig 1.**
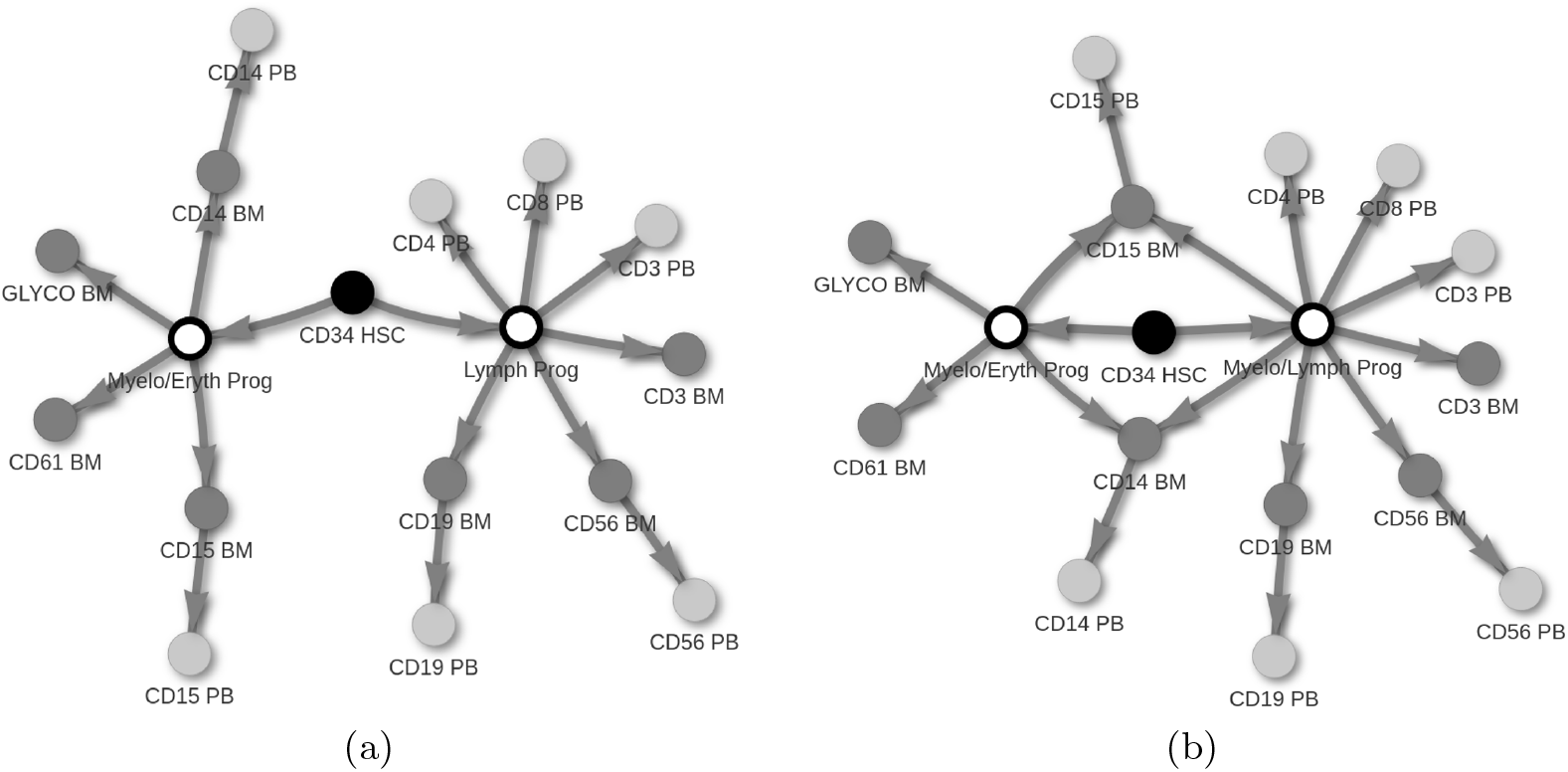
Two competing hemotopoiesis theories. Filled nodes correspond to lineages analyzed in this manuscript. Black, dark gray and light gray nodes represent Hematopoietic Stem Cell, Bone Marrow, and Peripheral Blood lineages. Empty nodes are latent, unobserved cell types. The classical dichotomy model (a) assumes that HSC first generate a common myeloid-erythroid progenitor (CMEP) and a common lymphoid progenitor (CLP), whereas the alternative myeloid-based model (b) postulates that HSC first diverge into the CMEP and a common myelo-lymphoid progenitor (CMLP).

This study aims to provide novel insights about human hematopoiesis and the HSC differentiation process in-vivo. This crucial biological question remained unresolved despite extensive efforts over the past years. Exploiting clonal tracking data from WAS gene therapy clinical trial, in section 6 we will investigate the hierarchical relationship among cell types and estimate lineage-specific cell duplication and death rates.

## 3. Stochastic logistic cell differentiation process

We consider an *N* -dimensional, continuous time counting process ***X***_*t*_ = (*X*_*t*1_, …, *X*_*tN*_), where *t* ∈ ℝ and 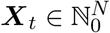. Each element of *X*_*ti*_, corresponds to the number of cells of type *C*_*i*_, (*i* = 1, …, *N*) present in the system at time *t. X*_1_ refers to the HSC count, the most primitive and multi-potent cell type.

We assume that ***X*** evolves according to a continuous-time Markov process. There are three event types in the process: cell duplication, cell death, and, importantly, cell differentiation. Individual cells are assumed to evolve independently from each other and cells belonging to the same cell type are assumed to obey the same laws. Event rates are assumed constant over time. The generic cell duplication rate *α*_*i*_ ≥ 0 is assumed to be a linear growth term, corresponding to the expected number of cell duplications per time unit per cell of type *C*_*i*_, *i* = 1, …, *N*,

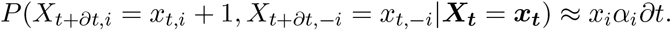

Secondly, linear cell duplication is eventually overcome by quadratic cell death. This assumption results in a cell type specific logistic growth curve, represented by the following conditional transition probabilities for cell death of type *C*_*i*_ (for some *δ*_*i*_ ≥ 0, *i* = 1, …, *N*),

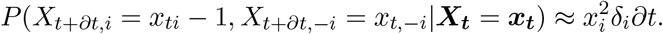

Furthermore, it is assumed that cell differentiation from cell type *i* into cell type *j* is a process with constant rate *λ*_*ij*_ ≥ 0, *i, j* = 1, …, *N, i* ≠ *j*,

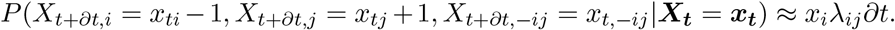

It is convenient to write the Markov process in a vectorized form. Each cellular event *k* ∈ {1, …, *r*} can be associated with an *N* -dimensional integer vector ***v***_*k*_, describing the net change in the state induced by event *k*. Given the hazard 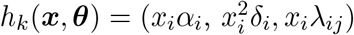 for ***θ*** = (***α, δ, λ***), we can write the process generally as *P* (*X*_*t*+*∂t*_ = *x*_*t*_ + ***v***_*k*_ | *X*_*t*_ = *x*_*t*_) ≈ *h*_*k*_(*x*_*t*_; *θ*)*∂t*. The whole process can be recast in matrix notation involving the net effect matrix, ***V***, corresponding to an *N* × *r* integer matrix, in which the columns correspond to the vectors ***v***_*k*_ (*k* = 1, …, *r*). For simplicity we assume that the first *N* columns of ***V*** refer to cell duplications, the second *N* to cell deaths and the remaining columns to differentiation events. The hazard ***h***(***X, θ***) = (*h*_1_, …, *h*_*r*_), is the *r*-dimensional vector of the *r* individual event hazards.

We are here considering that cells can only divide symmetrically, generating two daughters cells of the same nature as the mother cell. Assuming the alternative asymmetric division, such as in Xu et al. (2019), means that division is always coupled with a differentiation event, resulting in the formation of two cells with different properties and fate. Even though recent literature based on in-vitro experiments supports the possibility for HSC to undergo asymmetric division, little is known about the frequency of such events in-vivo and whether other lineages also have this capability.

Logistic differential equation models are widely used in the study of hematopoietic dynamics. Yet, it has not been applied in the context of clonal tracking data. According to the transition probabilities specified in our cell differentiation process, a clone will generate new cells purely based on its current counts. When a given size is reached, scarcity of nutrients and space in the niche makes cells die at a faster rate, preventing clone size from growing exponentially. Biologically, this is likely to be a too simplistic model of steady-state maintenance. In-vivo, cell duplication and death are regulated based on the current system needs using complex signaling mechanisms. However, our assumption has the remarkable advantage of allowing inference on all parameters of the differentiation process, avoiding the necessity to resort to literature data to set some coefficients, as proposed in Xu et al. (2019), or to infer *net rates* (duplication minus death rates) as done in Pellin et al. (2019).

### 3.1. Moment equations

For any stochastic process obeying the Markov property, given some initial condition ***X***_**0**_, it is possible to determine the evolution of the probability distribution function associated with the system states over time, *P* (***X***; *t*), using the Chemical Master Equation (CME) (Bailey, 1964; Kampen, 1981; Risken, 1984; Gardiner, 1985). The CME is defined as a differential equation for the process transition probabilities and can be written as

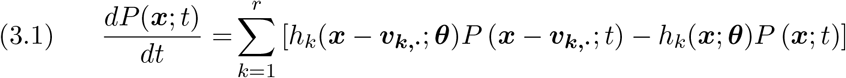

A solution and complete characterization of *P* (***x***; *t*) from (3.1) is unfeasible due to the large set of possible states configurations. However, important insights about cell differentiation dynamics and its parameters can be determined based on the time evolution of a few low-order statistical moments. Let *m*_*i*_(*t*) describe the time evolution of 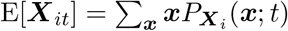. Applying the derivate operator to both sides, we obtain the dynamics of the mean of ***X***(*t*) can be summarized in the following ODE system,

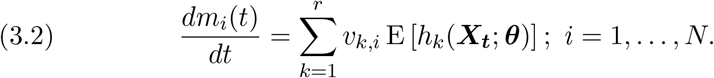

Similarly, let 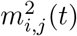 be the time evolution for the symmetric second-order moments E[*X*_*ti*_*X*_*tj*_] as

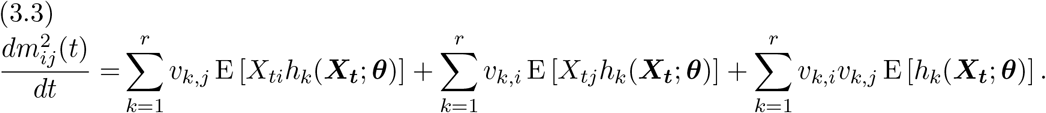

A detailed derivation of (3.2) and (3.3) can be found in Supplement A. With death rates 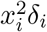 being polynomial of degree 2, the time evolution for the generic moment of order *n* depends on moments of order *n* + 1, leading to an infinite system of equations that can not be solved directly. There are different approaches to address this issue that consists of approximation methods. The most popular are the Chemical Langevin Equation, a diffusion approximation of the CME (Wilkinson, 2006; Golightly and Wilkinson, 2005), the system size expansion (Kampen, 1981; Elf and Ehrenberg, 2003), the Linear Noise Approximation (Gardiner, 1985), and the moments closure approximation (Grima, 2012). Hematopoietic differentiation is a stochastic process with an output consisting of a relatively small amount of cells, that starts from an individual HSC. These are not ideal conditions to apply the CLE approximation (Schnoerr, Sanguinetti and Grima, 2017). In its fundamental formulation, LNA requires the assumption that fluctuation and, as a consequence the clone cell counts, have a multivariate Normal distribution. This assumption, combined with the deterministic first-moment dynamics, poses challenges for approximating systems with multimodal steady-state behavior, as it is the cell differentiation process. We therefore approximate the moments evolution using moment closure.

Several moment closure approaches have been proposed in the literature: (i) assuming a specific probability distribution for *P* (***X***; *t*) (Whittle, 1957; Nåsell, 2003a,b; Keeling, 2000) or (ii) a separable-derivative-matching schema proposed in Singh and Hespanha (2007). The choice of the most appropriate method depends on the application of interest and the nature of the data analyzed. In this manuscript, we follow the indication provided in Schnoerr, Sanguinetti and Grima (2017), where these methods have been thoroughly tested and compared. Based on numerical evaluations, authors conclude that moment closure based on a normal distribution assumption is in general favorable for stability and precision. However, it is important to notice that the approach presented here is in principle valid irrespectively of the moment closure strategy adopted.

A Gaussian third-order moment approximation consists of setting the skewness equal to 0, leading to third-order moment definitions as follows,

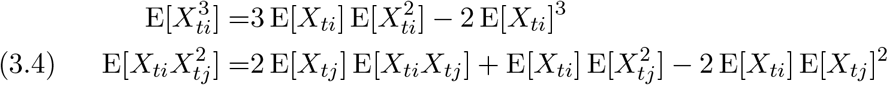

Substituting these third-order moments in (3.3) with the appropriate nonlinear formulation in (3.4), we derive two coupled systems of ordinary differential equations for the first and second order moments for the stochastic cell differentiation process. Based on this ODE system we will now propose an inferential procedure able to obtain parameter estimates and to reconstruct the cell differentiation structure.

## 4. Inference

The cell differentiation process is typically observed across a discrete number of time points and some replicates. To simplify notation, we assume we have *S* equally Δ*t*-spaced observations ***X***_***s***_, *s* = 1, … *S* from one realization of an *N* -dimensional stochastic cell differentiation process. It is computationally trivial to drop the equal spacing assumption. A likelihood-based approach would need to integrate all possible states and intermediate time-points, effectively making closed-form inference impossible. Instead, we will derive a methods-of-moments type estimator for inferring the parameters of interest.

As mentioned in section 2, in an experimental setting, clone sizes are estimated using NGS readouts. Despite several protocols, techniques and estimators proposed in the literature (Berry et al., 2012; Calabria et al., 2014; Leonardelli et al., 2016), measurement error still plays an important role in the quantitative characterization of the progeny of an individual HSC. Therefore, we included in our model definition (4.1) a multiplicative noise term that can be adjusted using an intensity parameter to be set according to the protocol followed.

### 4.1. Non-linear generalized method of moments

We reformulate the process as a non-linear regression problem, i.e.,

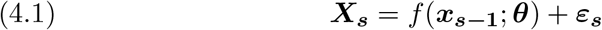

where *f* (***x***_***s*−1**_, ***θ***) = *E*[***X***_***s***_|***x***_***s*−1**_; ***θ***] is a known non-linear function of the process state at time step *s* − 1 and ***ε***_***s***_ is an *N* -dimensional mismatch variable with E[***ε***_***s***_] = **0**_***N***_, Var(***ε***_***s***_) = ***W***_***s***_ +*φ****N***_***s***_. ***W***_***s***_ = Cov[*X*_*i*_(*s*), *X*_*j*_(*s*)|***x***_*s*−1_; ***θ***] is a *N* × *N* matrix for some known non-linear function *g* modeling the stochastic process intrinsic covariance structure. The diagonal matrix *φ****N***_***s***_ describes a multiplicative-like noise term that allows to include a measurement uncertaninty on cell counts recordings. In particular, *φ* is a user-defined dispersion parameter that can be set by using a control experiment, as described in section 6, and ***N***_***s***_ = Diag(***x***_***s*−1**_) is a *N* ×*N* diagonal matrix with the cell counts on the diagonal. To avoid the usage of unnecessarily complicated notation in the description of our inference framework, throughout this section we will consider observations to be noise-free (*φ* = 0). However, the implemented method on the data does consider the dispersion parameter (*φ* > 0).

For each value of *s* the function *f* (***x***_***s*−1**_, ***θ***) = ***m***(*s*) and matrix ***W***_***s***_ = ***m***^2^(*s*) − ***m***(*s*)***m***(*s*)^*t*^ are defined through the solutions of the coupled ODE system (3.2) and (3.3) setting ***x***_***s*−1**_ as initial conditions for ***m***(*s* − 1) and *x*_*s*−1,*i*_*x*_*s*−1,*j*_ for 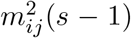. This projects the state and covariance matrix from one observed time-point to the next.

Applying this procedure to all observations available, we can perform parameter estimation by means of a generalized method of moments estimator with objective function,

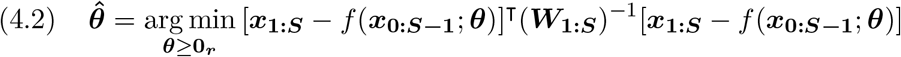

where ***x***_**1:*S***_ and *f* (***x***_**0:*S*−1**_; ***θ***) are (*N* × *S*)-dimensional column vectors and ***W***_**1:*S***_ is a *NS* × *NS* block diagonal matrix, in which blocks correspond to expected variance-covariance matrices ***W*** _*s*_ within each time increment. In Supplement B all the elements introduced in this section are derived for a simple example involving 3 cell types.

To calculate the solution 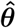, we propose an iterative procedure in which moments estimation and parameter refinement alternate until a convergence criterion is met. The complete algorithm is described in section 4.2. It is worth noting that for the solution of (4.2), some initial values 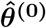for ***θ***, must be provided as input in order to start the iterative optimization procedure. Given the amount the parameters involved in the model, especially if no or limited assumptions are made to limit possible cell differentiations (by setting *λ*_*ij*_ = 0 for some *i, j*), it is important to start the minimization of (4.2) from accurate starting values within the convex region surrounding the true, unknown ***θ***. Supplement E presents a local linear approximation approach that can be used to obtain a sensible starting value (Pellin et al., 2019).

### 4.2. Algorithm

To find the solution to the minimization problem in (4.2), a modified implementation of the Gauss-Newton algorithm is proposed (Björck, 1996). Its pseudo-code is available in Algorithm 1. The procedure receives as input the initial cell counts, observations during the follow-up time, ***x***_**0:*S***_, and the system of ODEs for the first order, ***m***(*t*), and second order, ***m***^**2**^(*t*). The algorithm starts with the initial estimate 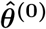 that is then refined using an iterative procedure with the updating formula

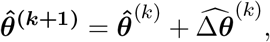

where 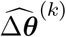 is the solution to the following constrained quadratic problem,

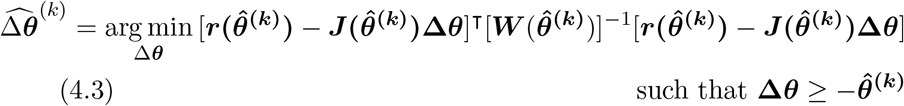

in which 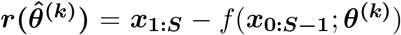is the residual *NS*-dimensional column vector and

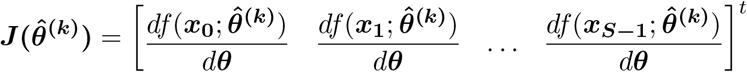

is the *NS*×*r* Jacobian matrix. Each 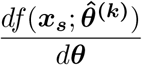 is a *N* ×*r* matrix measuring the change in predicted evolution for the mean of each component of the process caused by a small displacement of parameter vector around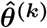. Finally,

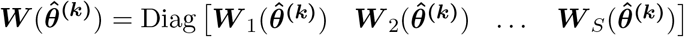

#### Algorithm 1

Iterative procedure for the non-linear generalized method of moments based parameter estimation.

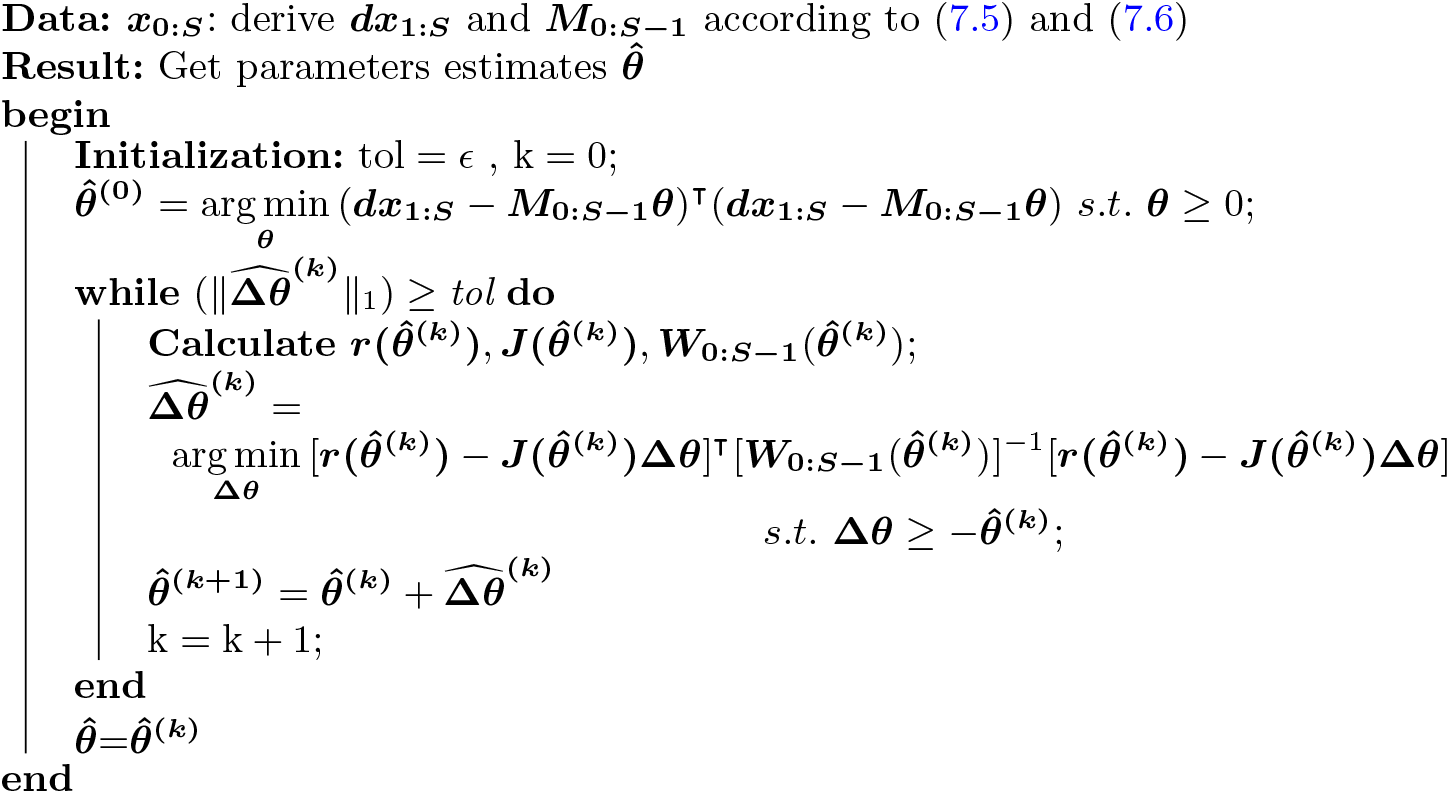

is the estimated *NS* × *NS* covariance matrix, setting the parameters vector to current value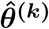.

For the local linear approximation method, some modifications to Algorithm 1 have to be made. At each iteration, parameter refinement is not performed by estimating increments vector 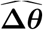, but 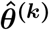 directly by solving the generalized (constrained) least square problem in (7.7) with covariance matrix calculated using 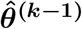.

## 5. Simulation study

In this section, we present four simulation studies. In the first, we study the behavior of the non-linear inference procedure simulating the data under that very model. In particular, we compare the method to a linear alternative, known as the local linear approximation, for several sampling intervals. For short sampling intervals, it is expected that the local linear approximation will be a serious competitor, whereas for longer sampling intervals the non-linearity will start to favor our non-linear inference scheme. In the second simulation study we mimic an experimental setting scenario by perturbing clones trajectory with multiplicative errors before performing inference. Our goal here is to investigate how an additional and extrinsic source of variation affect parameter estimation. The third simulation study focuses on how our model deals with model misspecification. Although our model is a detailed and generative model of the cell differentiation process, it is almost certain that this model is wrong — as all models are (Wit, Heuvel and Romeijn, 2012). We report the performance of our model in recovering the differentiation process under an alternative generative model. The fourth simulation study compares our proposal to an alternative method-of-moments formulation proposed by Xu et al. (2019), based on matching model-based and empirical correlations among cell types dynamics.

### 5.1. Improvement over local linear approximation approach

The inference procedure presented in this paper requires one to calculate as many solutions of the system of non-linear ODEs related to the first and second moments of the process, as available observations. In Figure 2a the network representation of the simulated system is shown. The precise parameter settings are given in supplementary materials Supplement D.

**Fig 2.**
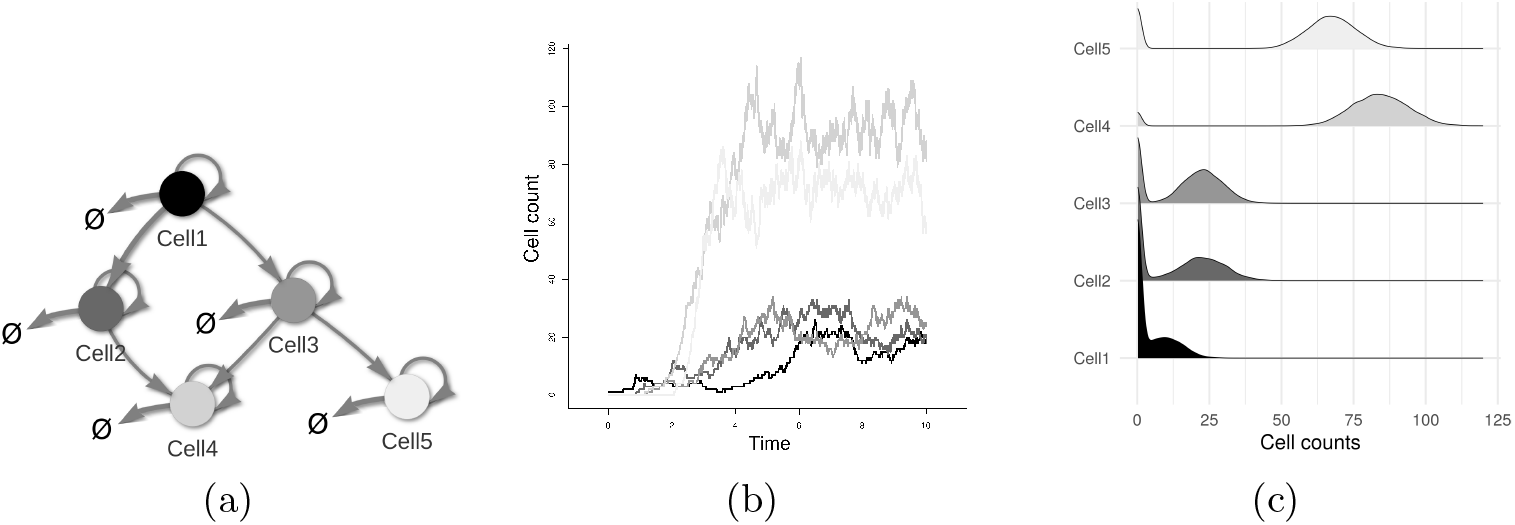
Details on the cell differentiation process used in the simulation study. A) Structure of the 5 cell types stochastic cell differentiation process. Cell types are represented by nodes. Self-connecting edges are duplication events. Death events are expressed by edges pointing to ø. Edges connecting two nodes correspond to differentiation paths. B) An example of cell differentiation process trajectory (clone evolution) generated by means of Gillespie algorithm. C) Cell types multi-modal steady-state distribution calculated using 1000 trajectories.

The stochastic cell differentiation process implemented has been designed with a low number of cell differentiations (5 out of 20) to reflect the expected scenario of real biological systems. The simulation study aims to determine whether our procedure is capable of correctly estimating the process parameters (both positive and zeros) and to investigate its performance for different sampling intervals. Clone dynamics are simulated employing the Gillespie algorithm (Gillespie, 1977), known to generate statistically correct trajectories of the stochastic equation described in (3.1). An illustrative trajectory is shown in Figure 2b, where it is possible to appreciate the logistic behavior generated by the model specification. Continuous-time trajectories are then sampled at three different equally spaced time intervals Δ*t* = (0.1, 0.5, 1) until stopping time *t*_end_ = 10 is reached. Parameter estimates obtained by using the proposed algorithm and the local linear approximation approach are compared for 100 experiments, each composed of *n* = 1000 clones starting from initial conditions vector ***x***_0_ = (1, 0, 0, 0, 0). Having clone evolutions starting from a single cell makes steady-state behavior particularly sensitive to the initial (stochastic) sequence of cellular events. In Figure 2c the distribution at *t*_end_, calculated based on 1000 clone trajectories, highlight the presence of a multi-modal steady-state configuration. On average, our algorithm converges in 2.8, 4.2 and 5.9 iterations, respectively, for Δ*t* equal to 0.1, 0.5 and 1. The local linear approximation approach converged on average in 3.2, 6.2 and 7.2 iterations.

As shown in Figure 3, the local linear approximation based method suffers in terms of accuracy in all settings. Due to the limited amount of cells present in the system in the initial phases, the strong non-linearity component of the dynamics is poorly approximated by the linear approach. As a consequence, there is a considerable and fast decay of estimation precision as Δ*t* increases. For Δ*t* = 0.1, the local linear approximation approach seems to be able to recognize the underlying structure of the system, since almost all absent differentiation paths are correctly estimated as very closed to zero. This is not true for larger time gaps Δ*t*, e.g., 0.5 or 1, where, in addition to a considerable bias for all estimates, some of the absent links – for example, *λ*_1,5_ is shown in Figure 3 (bottom-right) – are systematically estimated as greater than 0. The non-linear inference procedure, instead, shows unbiased estimates for all Δ*t* considered for all parameters.

**Fig 3.**
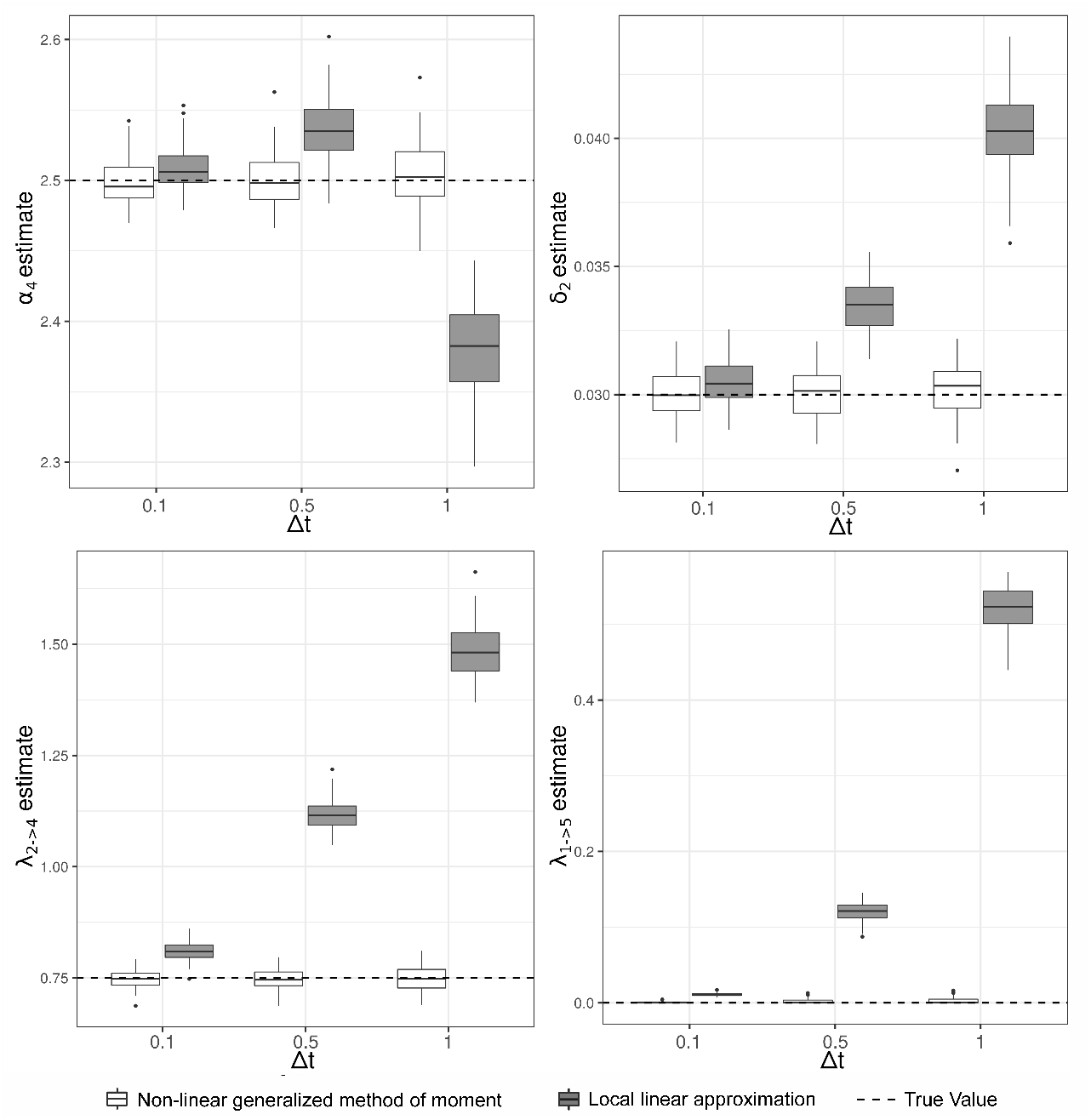
Comparison between the non-linear generalized method of moments and local linear approximation for different Δt setting. Estimate distributions for the non-linear generalized method of moments and the local linear approximation are displayed using respectively white and gray boxplots. Dashed lines correspond to the true values. On top-left the performance of the methods for the estimation of the duplication rate α4 (2.5) is shown. On top-right, death rate δ2 (0.03) is reported. On the bottom the differentiation rates λ1,3 (0.35, left) and λ1,5 (0, right) are represented.

### 5.2. Performance introducing measurement errors

To investigate how measurement errors affect the performance of our proposal for inference, we apply our algorithm to perturbed clone trajectories, 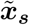. These trajectories are generated by adding noise to the exact one, ***X***_*s*_, as follows

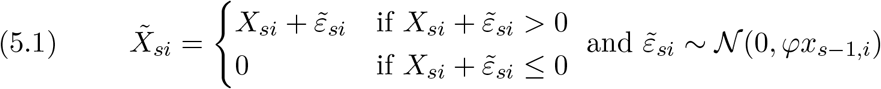

We considered the same system configuration and experiment setup as described in section 5.1, using Δ*t* = 1 and inspecting the impact of noise of different strength by testing *φ* = (0, 0.1, 0.5, 1).

In Figure 4 the performance in estimating a duplication rate (*α*_4_), death rate (*δ*_2_), differentiation rate (*λ*_2,4_) and an absent differentiation path (*λ*_1,5_) is shown. For all parameters, we observed an increase in the standard errors as the value of *φ* increases. A shift in the parameter distribution is observed for death and differentiation rates for the larger values of *φ*, but not for the duplication coefficient. Most likely for large values of *φ*, as the states are artificially truncated at 0, probably a bias is introduced. The higher *λ*_1,5_ average estimates we observed for *φ* values (0.5,1) is presumably due to the increase of the estimator standard error. The vast majority of *λ*_1,5_ estimates fall in the (0,6e-4) range, suggesting that the correct identification of missing differentiation paths is robust to higher levels of observational errors. To recover the underlying network structure and eliminate potential spurious, low-intensity connections among lineages, in section 6 we propose a model selection strategy based on backward stepwise selection and cross-validation.

**Fig 4.**
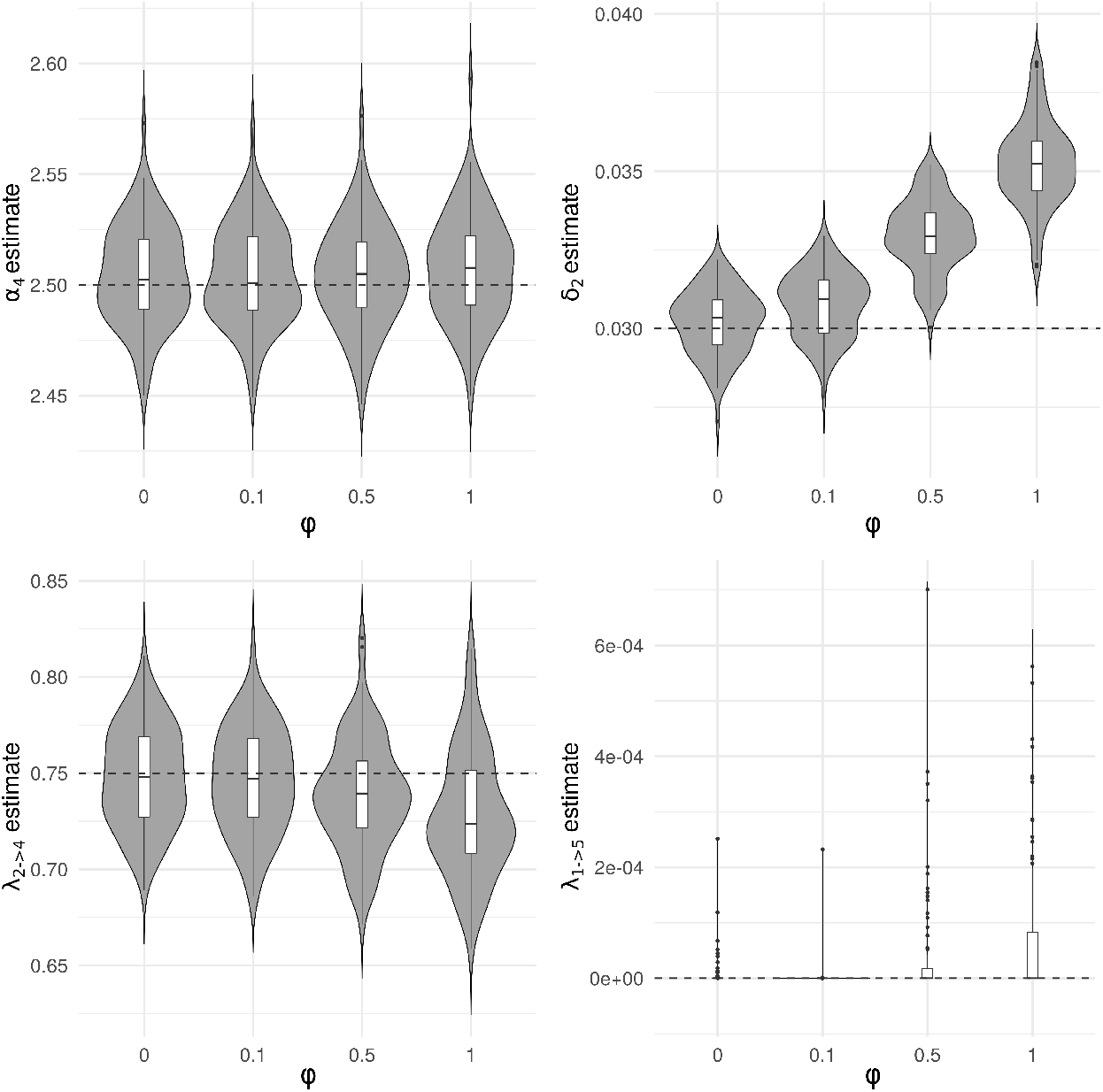
Impact of measurement errors on inference. On top-left the performance of methods for duplication rate α4 (2.5) estimation. On top-right the performance of methods for death rate δ2 (0.03) estimation. On bottom-left the performance of methods for differentiation rate λ2,4 (0.75) estimation. On bottom-right the performance of methods for absent differentiation path λ1,5 (0) estimation.

### 5.3. Performance under model misspecification

There are various difficulties associated with modeling biological processes, in particular when dealing with questions related to the *in-vivo, in human*, investigation of complex phenomena such as hematopoiesis. Many reasons limit sample size and the type of experiments that can be performed, forcing the researcher in making important assumptions about biological mechanisms based on evidence gathered from *in-vitro* or animal studies, not always representative of human dynamics. For these motivations, it is important to check how new statistical procedures behave in case of model misspecification. In order to test our proposal described in section 4 under this condition, we generated clone trajectories using a corrupted version of the Gillespie algorithm. Differentiation process structure has been kept as shown in Figure 2a. Parameters have been set to the same values as reported in section 5.1, except for death rates set to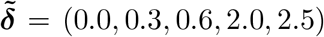. Individual clone evolution has been simulated as described in the following steps:

1. Set initial state at ***x***_0_ = (1, 0, 0, 0, 0).
2. Generate time-to-next event, *t*_*s*+1_, sampling from a Uniform distribution with parameters Unif 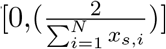.
3. Select a cell type, *C*_*s*+1,*i*_ sampling with probability proportional to cell count among those cell type with *C*_*s,i*_ ≥ 1.
4. Sample a cell event (duplication, death or differentiation) among those available for the specific cell type *C*_*s*+1,*i*_ with probability proportional to event rates.
5. If total event time is less than 10, return to step 2.

It is worth noting that these modifications affect multiple aspects of the data generating process, as visible from Figure 5a. Events frequency is much lower throughout the simulation period and cell counts do not stabilize around a cell type-specific value, as was the case for the original model shown in Figure 2b, but they rather exhibit exponential growth dynamics. In the correct version of the Gillespie algorithm, the time-to-next-event is distributed as an exponential with parameter exp 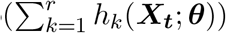 and the same vector of events hazard *h*_*k*_(***X***_***t***_; ***θ***) is rescaled to the unit sum in order to define events sampling probabilities. Under the misspecification setting, the event times are distributed uniformly, and the event probabilities are not directly linked to the hazards.

**Fig 5.**
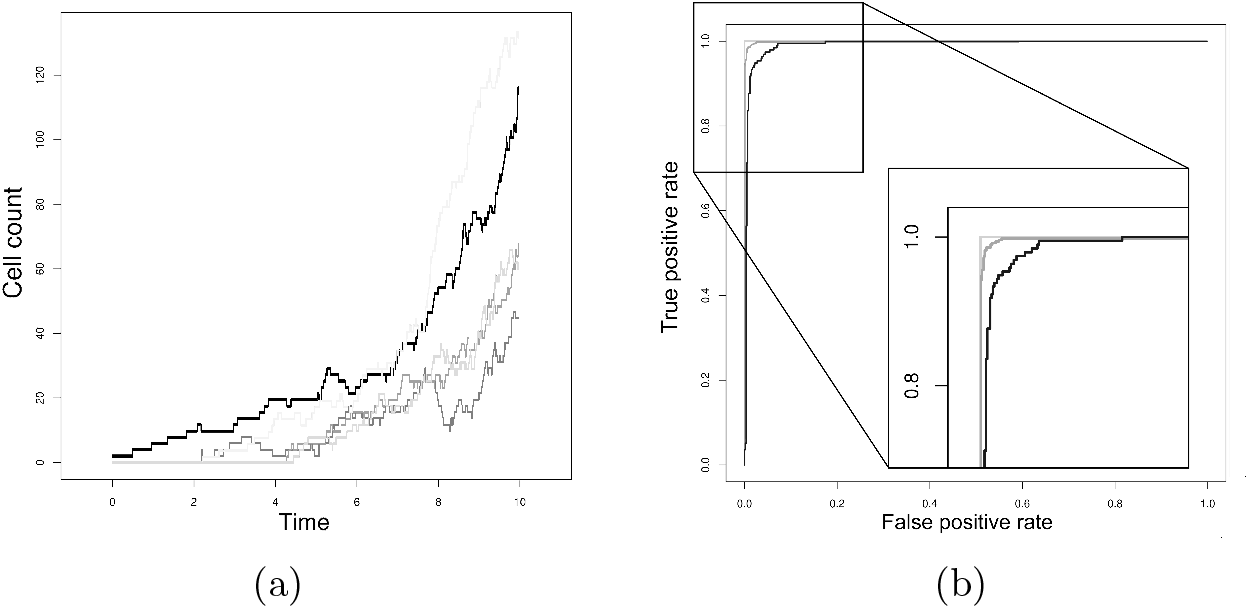
Misspecified cell differentiation process. A) Cell differentiation process trajectory generated by means of the misspecified generative model. B) Average ROC curves obtained from 100 experiments replicates each containing 30 (black), 50 (dark gray) and 100 (light gray) clones.

Three different sample sizes have been tested: 30, 50, and 100 clones per experiment. We evaluate our inference method for its capability to correctly reconstruct the underlying differentiation structure, rather than for the precision in parameters estimation. Based on the data generated from a single experiment, we test the null hypothesis *H*_0_ : *λ*_*ij*_ = 0, *i, j* = 1, …, *N, i* /= *j* as described in Supplement C.

Each ROC curve in Figure 5b shows the average of 100 ROC curves obtained from independent replicates of the simulation experiments by varying the significance threshold on differentiation rates. Our generalized method of moments approach shows surprising accuracy in learning the true network configuration for 30, 50, and 100 clone trajectories for a wide range of significance threshold values.

### 5.4. Comparison with correlation-based M-estimator by Xu et al. (2019)

Although the general model specification of the clonal expansion dynamics by means of stochastic differential equations is similar to that described in Xu et al. (2019), there are several differences between the data-handling and estimation approaches. Wu et al. (2014) only have clone size measurements in 5 mature blood lineages and no information on the progenitors. To estimate the hidden relationships among stem and progenitors cells, Xu et al. (2019) resort to comparing known tree-like differentiation configurations by means of cross-validation. Furthermore, in order to obtain an analytical solution for the moments evolutions, they assume event hazards to be linear in process states. This is probably the only sensible workable assumption, but it does imply either exponential extinction and growth dynamics of the clones. On the other hand, the gene therapy study motivating our method consists of 15 cell types from both BM and PB, providing a much more detailed description of the complete hematopoietic process. Given this motivation, we designed a modeling approach that assumes all lineages of interest to be observed.

In order to compare the two methods, we modified our methodology to consider, as in Xu et al. (2019), asymmetric division (differentiation is coupled to cell division) rather than symmetric division, whereby cell duplication is followed by a differentiation event. Furthermore, to match the two stochastic processes we assumed that the dynamics does not involve saturation by assuming linear ODEs. To make a reasonable comparison among the two methods under the fully observed scenario, we extended the calculation of correlation-based M-estimator proposed by Xu et al. (2019) to all correlations among lineages, including stem cells, progenitors and mature cell types.

We set up a simulation study resembling the one described in Figure 2c in Xu et al. (2019) (reproduced here in Figure 6a) both in terms of the differentiation tree structure and the rate parameters. The process consists of 8 cell types, starting from a *HSC* that duplicates wit rate *λ* = 0.285 and differentiates in progenitor cells, *Prog A* and *Prog B*, with rates *ν*_*a*_ = 0.14 and *ν*_*b*_ = 0.07, respectively. Progenitor cell-types A die with rate *µ*_*a*_ = 0.14 and differentiate into three mature cell types with rates *ν*_1_ = 36, *ν*_2_ = 18 and *ν*_3_ = 10, respectively. Progenitor cell-types B have two connected mature lineages into which it differentiates with rates *ν*_4_ = 20 and *ν*_5_ = 12. As done in Xu et al. (2019), we considered mature cells death rates known and equals to *µ*_1_ = 0.26, *µ*_2_ = 0.13, *µ*_3_ = 0.11, *µ*_4_ = 0.16 and *µ*_5_ = 0.09. All trajectories start with a single HSC at time *t*_start_ = 0. Each simulation experiment is composed of 1000 clones, observed at intervals Δ*t* = 1 unit apart, from *t*_start_ = 0 up to the final time-point set at *t*_end_ = 10. The results of the simulation study and the distributions of the coefficient estimates across 100 simulations are shown in Figure 6b.

**Fig 6.**
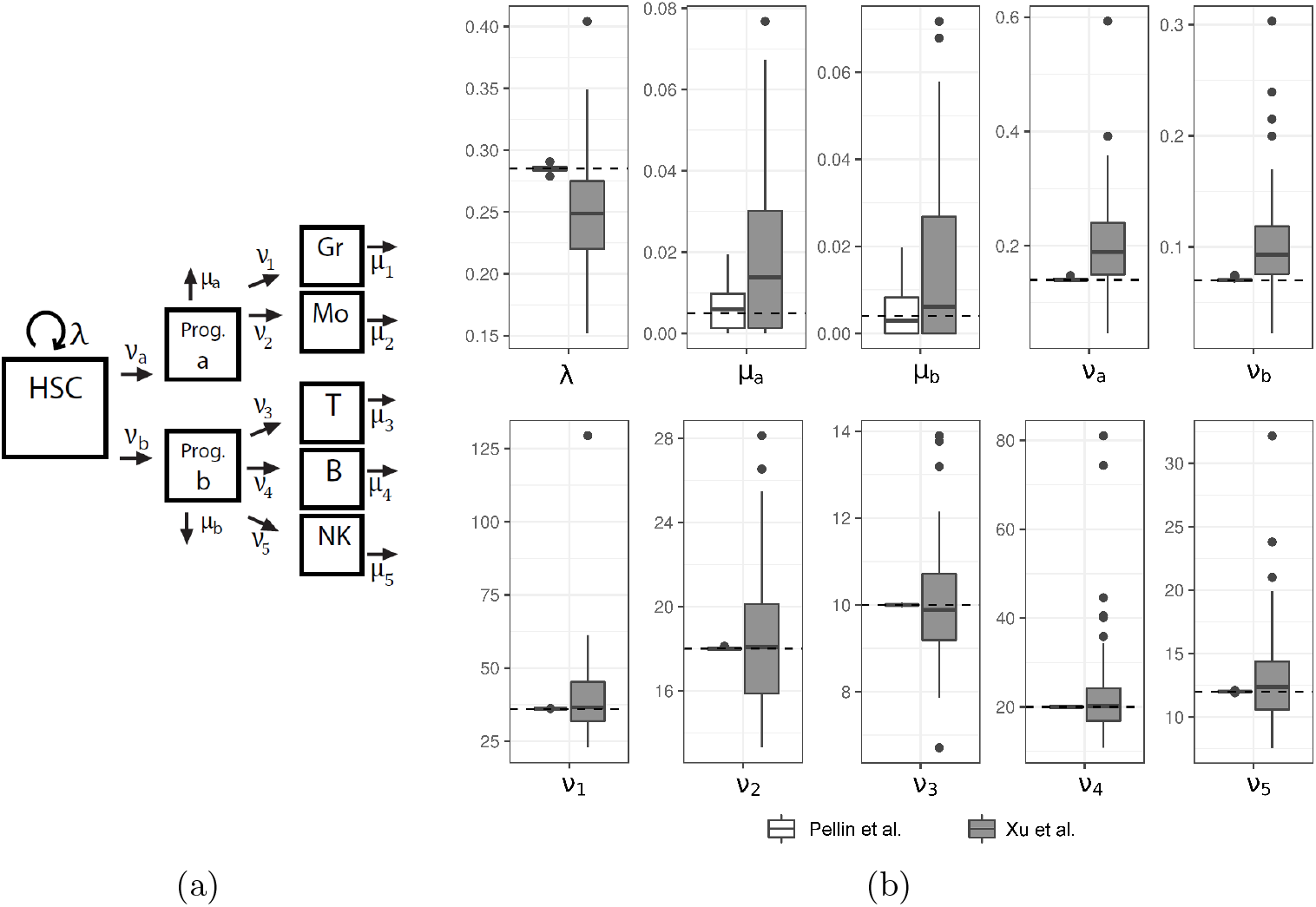
Comparison with Xu et al. (2019) correlation-based M-estimator. Considering the cell differentiation process shown (a), the boxplots in (b) show that our method is unbiased and more efficient than the M-estimator proposed in Xu et al. (2019). Boxplots show the distribution of estimates obtained using the method proposed in this manuscript (white) and the extended version of the correlation-based estimator (dark gray) for the 10 unknown rates, whose true value is indicated by the horizontal red dashed line.

Our proposal outperformed the method of Xu et al. (2019) in several aspects. The precision of our estimates is an order of magnitude better, and the bias of our method is negligible, whereas their estimation of *µ*_*a*_, *µ*_*b*_, *ν*_*a*_ and *ν*_*b*_ clearly suffers from bias. Furthermore, our computational algorithm converged in 4.3 iterations on average, whereas the correlation-matching algorithm converges on average in 60.6 iterations. The reason why our method outperforms the method proposed by Xu et al. (2019) is that latter based on second moment matching, whereas our method is based on first moment matching, which is more stable, unbiased and computationally more efficient. On the other hand, the main advantage of the method proposed by Xu et al. (2019) is that their method can deal efficiently with missing progenitor and HSC data. In certain experimental settings this can be crucial.

## 6. Gene therapy study for Wiskott-Aldrich Syndrome

In this section, we return to the previously described clinical trial treating patients suffering from Wiskott-Aldrich Syndrome with their stem cells, genetically modified ex vivo, and then reinfused to the patient. We traced *N* = 15 cell types over time in the three patients up to 36 months after GT. In Figure 7a the differentiation trajectories observed for two clones are shown. The 15 distinct cell types can be organized in a three levels hierarchy, corresponding to the original *HSC level*, i.e., CD34 stem cells, the *bone marrow (BM) level*, corresponding to CD3, CD14, CD15, CD19, CD56, CD61 and GLYCO precursor cells and finally the *peripheral blood (PB) level*, i.e., CD3, CD4, CD8, CD14, CD15, CD19 and CD56 mature cells. Based on the available biological knowledge, the following assumptions are made,

**Fig 7.**
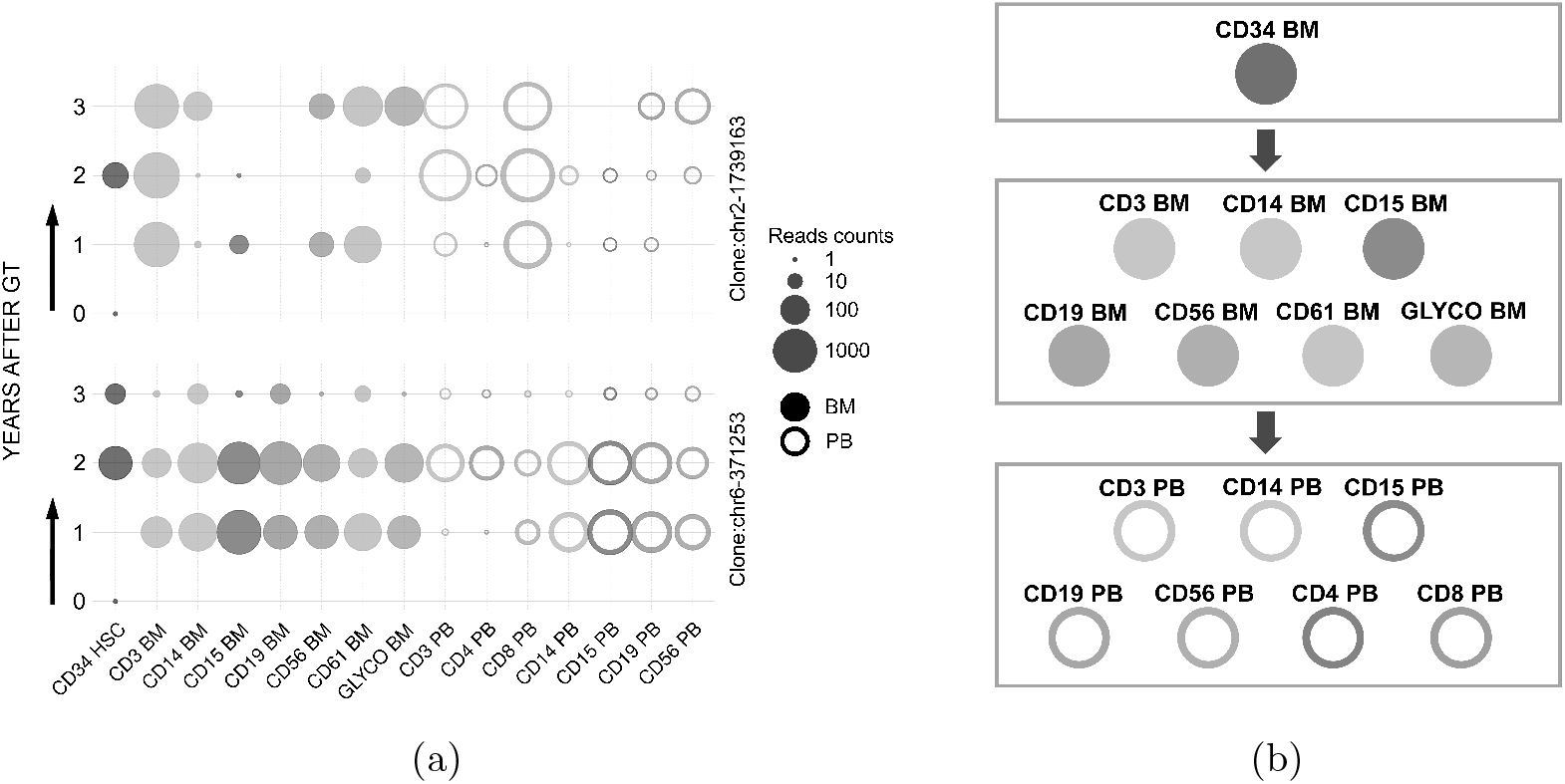
Observed clones dynamics and schematic representation of hierarchical assumptions. A) Reads counts trajectory over the 3 years follow-up for 2 clones. By assumption, all clones start with a count of 1 in CD34 HSC cell type at time 0. Cell count and are represented by circle of size proportional to their abundance. B) Schema reporting the biologically inspired three levels hierarchy used as a backbone for differentiation structure reconstruction. Arrows show directionality for potential differentiation paths.

- the HSC type can differentiate in any cell type in the BM level;
- cell types at the BM level can differentiate in any cell type in the PB level;
- cell types at the PB level can not differentiate.

These assumptions are graphically summarized in Figure 7b and incorporated in the stochastic cell differentiation model and inferential algorithm by setting the corresponding *λ*_*ij*_ to zero.

From a practical perspective, the re-infusion of corrected HSC cells in a patient’s body is considered as starting time *t* = 0. Initial conditions vector ***X***_0_ consists of a 15-dimensional vector, with the count corresponding to CD34, the HSC, equal to 1 and the rest to zero. During the follow-up period, *S* = 3 samples from patient’s HSC, BM, and PB cells are taken after 1, 2, and 3 years. After exluding all clones detected only once throughout the study period, in total we obtain 17,195 unique chromosomal positions: 5,299 from period 1, 5,300 from period 2, and 6,596 from period 3. The amount of cells, within each lineage, generated by individual labeled, re-infused HSC, is counted through an insertion site analysis technique described in Aiuti et al. (2013). For estimating the measurement error scaling coefficient associated with the protocol used in the processing of patients’ samples, we took advantage of the three independent experiments in which a pool of HSC cells have been sequenced 1-day after transduction. Given the low proliferative rate of HSC in culture conditions, all clones are expected to have a size of 1 at time of sequencing. Based on these data we estimated 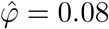.

### 6.1. Cell differentiation reconstruction

Clonal tracking studies typically score and compare alternative but fixed models of hematopoiesis using experimental data. In this work, we opted for data-driven learning of the differentiation process structure. To recover the actual underlying data generating process and eliminate differentiation paths caused by sampling issues and observational errors, we proceed as following described. We estimated the full model, *m*_0_, by solving the optimization problem 4.2 using the WAS data and 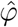. We then iteratively eliminate the differentiation connection (*λ*_*ij*_) with the least impact on the following Mahalanobis distance:

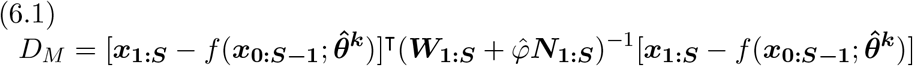

This method leads to a sequence of models, *m*_*k*_, *k* = 1 … 56 with decreasing complexity. To select the optimal model 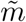 among the set *m*_*k*_, we used a 5-fold cross-validation strategy. We split the input dataset into five subsets of equal size and used four subsets to estimate the process parameters and the remaining as a validation subset on which the Mahalanobis distance (6.1) has been calculated. The procedure has been repeated five times for each model configuration, considering each subset for validation once. The results are reported in Figure 8. We selected model *m*_35_ as optimal based on its mimimum median Mahalanobis distance across folds.

**Fig 8.**
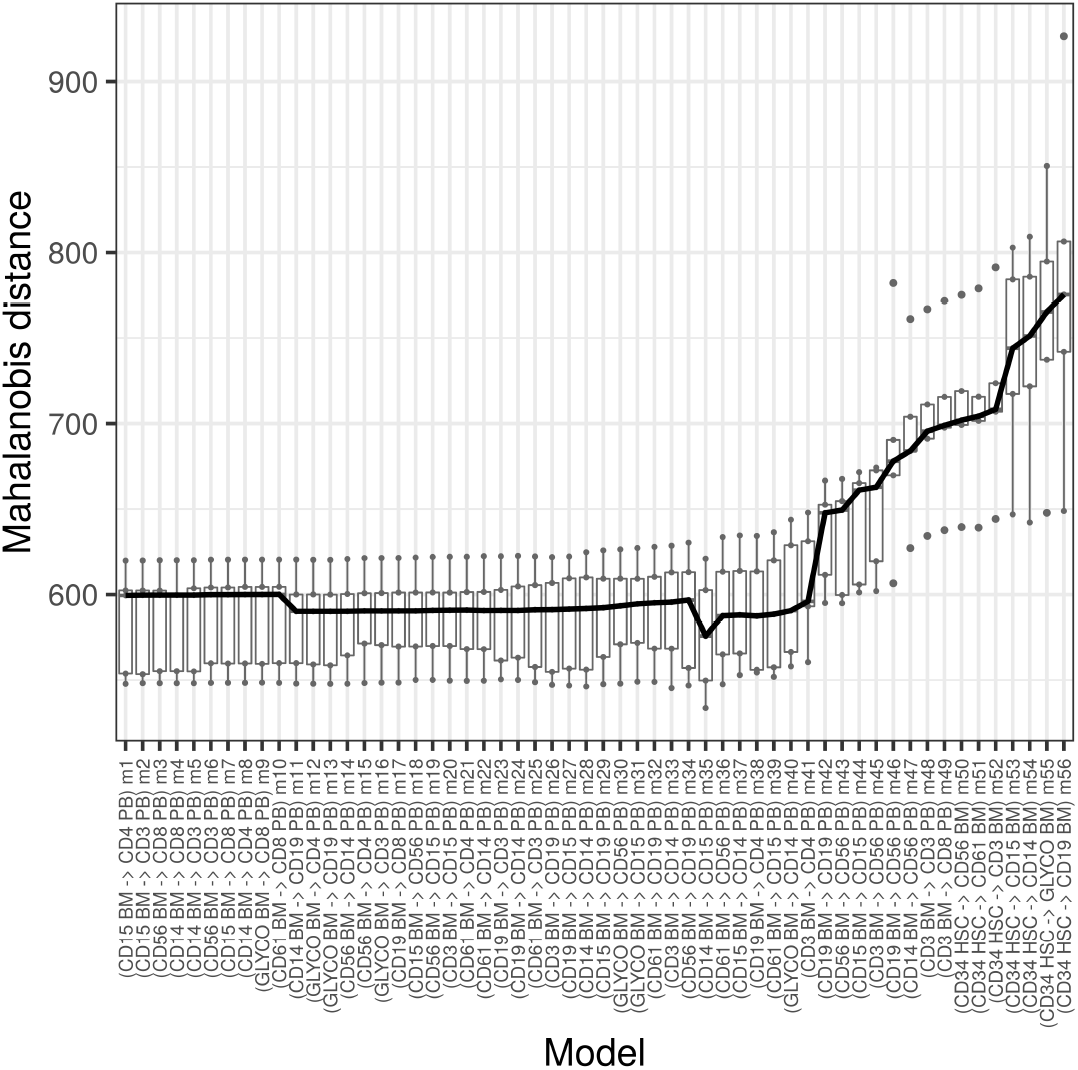
Cross-validation and Mahalanobis distance based selection of the optimal model. Models m_k_ are ordered from the most complex, m_1_ (one differentiation path removed from the full model m0) to the least complex, m56 (no connection among lineages). The additional differentiation path that is removed at each iteration is reported alongside model number. The distributions of the Mahalanobis distances calculated on the 5 validation subsets are represented with boxplots for each model configuration, m_k_. Solid black line connects the median distances across models. The minimum median is osberved for m35 that is therefore selected as the optimal model, 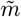.

We then imposed the differentiation structure encoded in model *m*_35_ and estimates the cell differntiation process parameters using all WAS data available. A graphical representation of the differentiation network is shown in Figure 9a. Duplication and death have been omitted in the plot for clarity, but all final parameters are available in supplementary materials Supple- ment F. In Figure 9b a trajectory of the HSC differentiation process estimated using WAS gene therapy data is shown, generated using the Gillespie algorithm.

**Fig 9.**
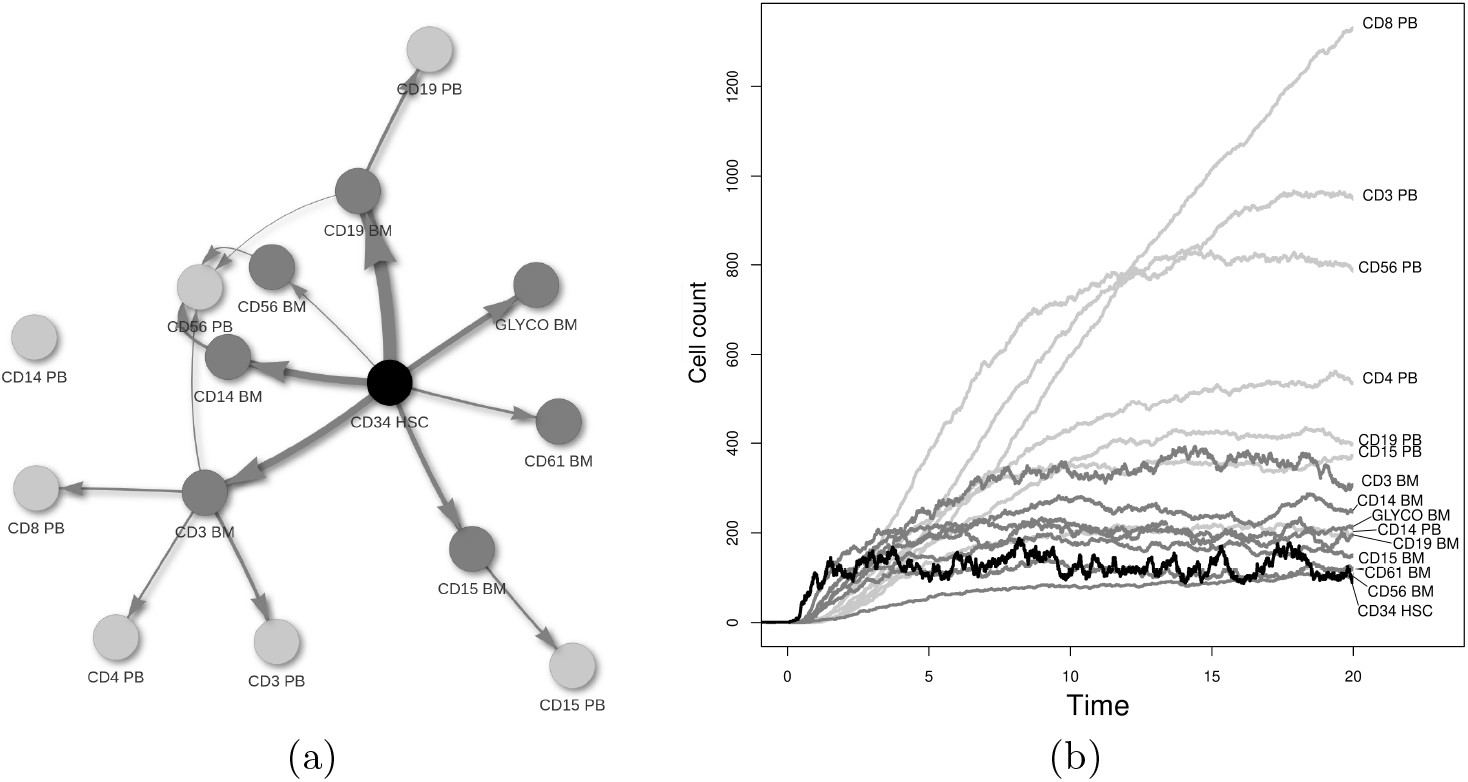
HSC differentiation process. (a) Network representation. Black, dark gray and light gray nodes represent CD34 HSC, BM, and PB cell types repsectively. Edge thickness is proportional to the corresponding 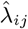. Edges estimated are those included in the optimal model 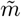. (b) HSC differentiation process trajectory simulated using the Gillespie algorithm, assuming model 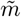 and coefficient estimated using WAS data.

**Fig 10.**
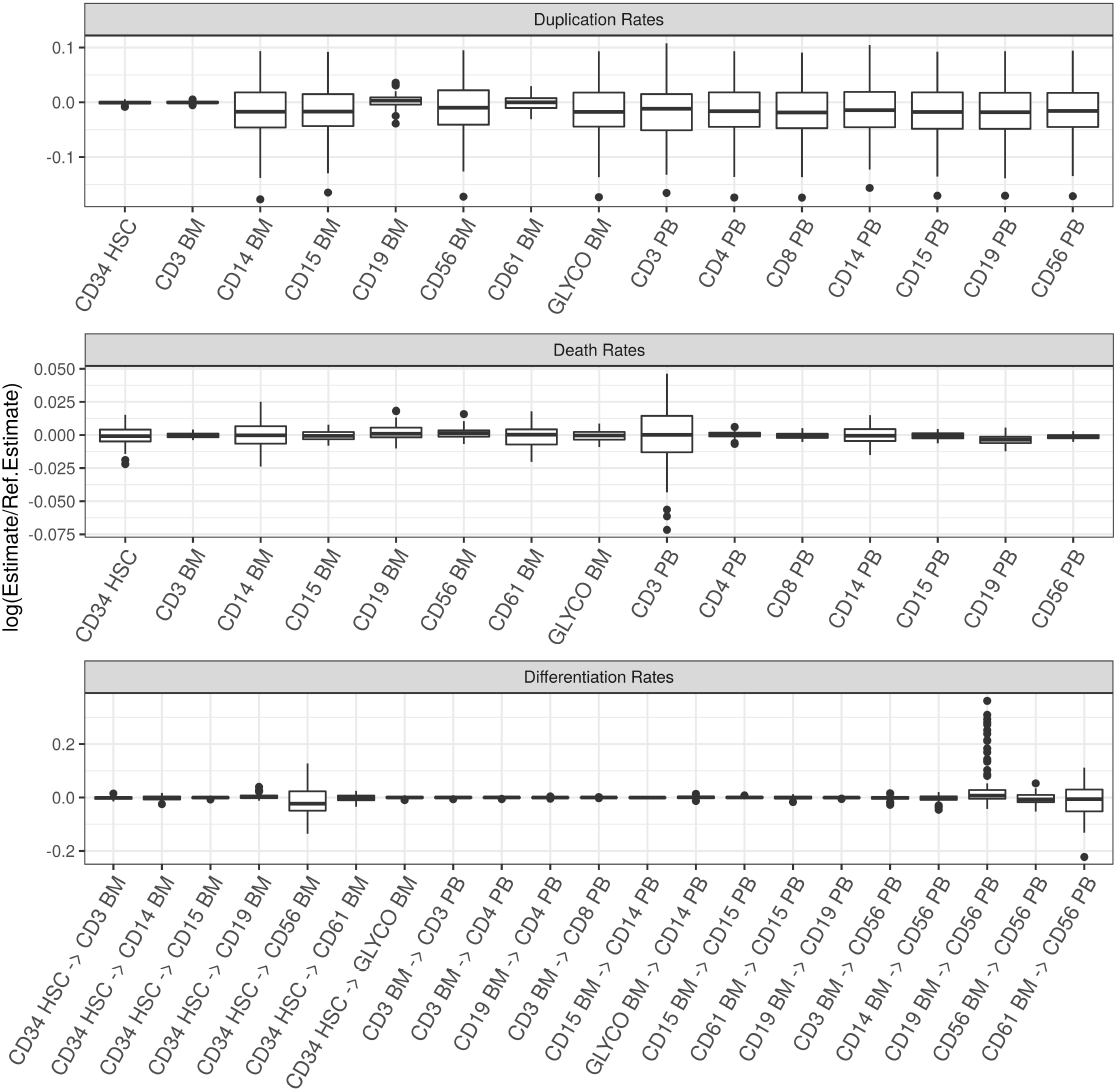
Sensitivity analysis to random initialization. Model 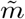 has been estimated starting from 100 different 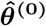 settings. Duplication and differentiation rates are sampled from a Normal distribution with 𝒩 (0.1, 0.1) and death rates from a 𝒩 (0.01, 0.01). Absolute value transformation was applied to avoid negative initial values. The distribution of logarithm of the ratio between the random restart estimates and the local linear initialization estimate (Ref. estimate, see values in Appendix Supplement F) is represented using boxplots.

Initialization with the local linear approximation aims at starting the optimization procedure in the proximity of the objective function global optima and reducing the number of iterations (*m*_35_ converges in 5 iterations) required to meet the convergence criteria. We verified that the parameters estimate in Appendix Supplement G are stable to random initialization by sampling candidate values for 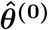 from a Normal distribution N (0.1, 0.1) for duplication and differentiation rates and N (0.01, 0.01) for death rates. We performed 100 random restarts showing that our estimates are robust.

### 6.2. Relevance of the results

CD34 HSC resulted in being the lineage with the highest duplication rate. According to our estimate, a CD34 HSC cell is expected to duplicate approximately every 6.51 weeks (*α*_*CD*34 *HSC*_ = 8.006*e* + 00), a significantly higher rate than the 40 weeks (range, 25-50 weeks) previously reported (Catlin et al., 2011). The difference is attributable to the following considerations. First, the patients enrolled in a GT clinical trial receive a conditioning regimen before treatment. Upon reinfusion, the transduced cells are subjected to high proliferative stress because they must replenish the depleted hematopoietic system. The estimate reported in Catlin et al. (2011) instead is referred to a healthy, native, steady-state condition and does not consider potential selective advantages that engineered cells might have in disease settings. Second, the CD34 marker used in the WAS study to isolate HSC from patients’ BM samples is known to select for a broader cell population that includes hematopoietic progenitors cells in addition to stem cells, which are characterized by a higher proliferative output and shorter half-lives compared to pure hematopoietic stem cells.

BM lymphoid lineages CD3 and CD19 show higher duplication coefficients than myeloid cell types (CD14 and CD15). This result supports the idea of the presence of long-lived lymphoid progenitors and the dependence of the myeloid compartment from the continuous support of cells coming from the upstream CD34 HSC population (see supplementary matierals Supplement G). CD61 BM cells are estimated to have a significant duplication rate. The distinct behavior of the megakaryocyte (CD61 BM) population is not surprising since megakaryo/erythrocyte-restricted progenitor, responsible for the production of platelets and red blood cells (erythrocytes), have been reported and validated in several studies, mostly based on gene expression data. Steady-state cell counts for individual lineages are not deterministic but depend on the specific evolution of each clone (see Figure 2c). However, in Figure 9b it is possible to appreciate how the combination of duplication and death rates estimate leads to a biologically meaningful differentiation process in which PB lineages are the most abundant, followed by BM and CD34 HSC.

In the optimal model configuration determined by our model selection strategy (Figure 9a), all BM lineages result directly connected to the HSC compartment. Surprisingly, HSC to B-Cell precursor (*λ*_*CD*34 *HSC*→*CD*19 *BM*_ = 1.453) differentiation rate is higher than HSC to myeloid cells (CD15 BM, CD14 BM), which are among the cell type with the fastest turnover in humans (Sender and Milo, 2021). This finding agrees with the conclusion of Meyer-Bahlburg et al. (2008) who, using mouse models of WAS, highlighted that upon transplantation, corrected B cells exhibit a marked selective advantage at both the precursor and mature stage.

The biology behind the maturation and migration of BM cells in the PB stream is much better understood, and commitment paths are well characterized. The consistency of our inferred structure at the BM and PB interface with the biological expectation is remarkable, even though a limited set of constraints to the network configurations has been provided. The separation between lymphoid and myeloid branches is clear, with significant differentiation parameters connecting CD3 at BM level to CD3-CD4-CD8 (T-cell) and CD56 (NK) in the PB. Among the myeloid subpopulations, CD15 BM is linked to CD15 PB as expected, but the differentiation from CD14 BM to CD14 PB is missing. The isolation of CD14 PB from all BM lineages is most likely a sampling issue since monocyte (CD14) account, on average, for only 5% of the cells in a PB sample.

Our results support the myeloid-based model over the classical dichotomy model. Mature NK cells (CD56 PB) are sustained by a cellular influx from NK cells residing in the BM (CD56 BM), as expected, but also from CD14 BM (myeloid), CD19 BM, and CD3 BM (lymphoid lineages). Although it is biologically challenging to conclude that all these cell populations can directly give rise to CD56 PB cells, this pattern is compatible with the presence of a common, unobserved progenitor cell type capable of generating both myeloid and lymphoid mature cells.

Due to the poor approximation provided by the local linear method, as also shown in our simulation study, Biasco et al. (2016) identified many more low-intensity, most likely spurious, differentiation rates. For this reason, the authors preferred to limit the inferential goal to calculate and compare the likelihoods of only two known and competing tree configurations using information-based criteria. Instead, the method presented in this paper allows us to perform network and coefficients estimation simultaneously. It requires only limited prior knowledge and is essentially data-driven. Nevertheless, it also offers the flexibility to trade exploratory power for biological interpretability by changing the settings of the differentiation rates fixed at zero according to the scientific question.

Finally, to resolve the conundrum regarding *in-vivo* stem cell evolution and hematopoietic differentiation structure, a more refined sorting strategy for HSC (CD34 BM) is needed. Through additional known surface markers, indicators of stem/progenitor cells priming towards specific lineages would be possible to disentangle the complexity observed at the BM level.

## 7. Conclusion

To improve our knowledge about the cell differentiation process, which in many contexts such as gene therapy might be fundamental for providing biological and therapeutic new insights, we have devised and implemented a flexible statistical framework for the analysis of clonal tracking data. The underlying stochastic process is assumed to be a multidimensional Markov process and this allows a representation of the moment dynamics by means of a system of non-linear ODEs. The particular definition of the transition probabilities induces a logistic behavior of sub-population growth curves. The model and the proposed iterative inferential procedure exhibit stability in terms of parameter estimation, structure recognition, and convergence rate. The model can easily be extended to incorporate time-dependent individual cell rates, different feedback regulation mechanisms, or random effects on specific parameters.

Applying the modeling and inference framework to a Wiskott-Aldrich Syndrome gene therapy study, we have obtained insight into the underlying stem cell differentiation dynamics. We found a high degree of agreement between our results and the recently proposed myeloid-based model for human hematopoiesis *over* the predominant classical dichotomy model of cell evolution.

## Acknowledgements

CdS and EW would like to acknowledge networking support by the COST Action COSTNET (CA15109). EW acknowledges funding from Swiss National Science Foundation (SNF grant: 188534).

## SUPPLEMENTARY MATERIAL

### Supplement A: Derivation of moments equations

(http://www.e-publications.org/ims/support/dowload/imsart-ims.zip). By means of the summation operator, 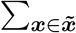, over the whole set of possible states for the process 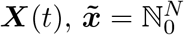, it is possible to derived a functional connection between the evolution for the expected population size of each process component and the dynamics of the process probability distribution *P* (***X***; *t*),

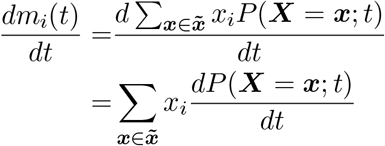

The evolution of *P* (***X***; *t*) can be expressed by means of the master equation introduced in (3.1),

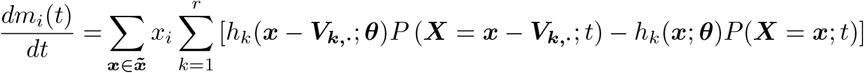

Due to the fact that the summation operator 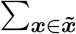 spans over all possible state configurations, the order of summation operators in the RHS can be inverted,

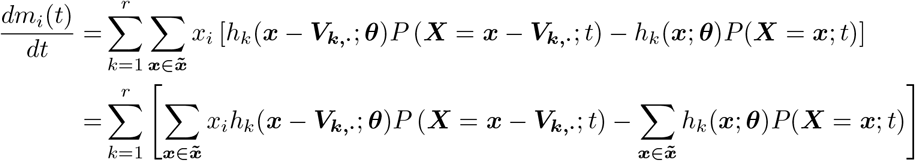

Now, the summation variable in the first term of the right-end-side can be modified, without affecting the sum domain, since it cover all possible state configurations,

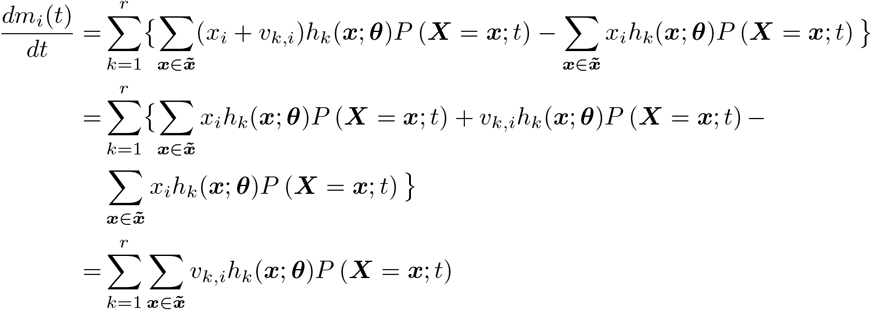

Given the known property for expected value of function *f* (*x*) of a r.v. *x* with probability distribution *P* (*x*), E[*f* (*x*)] = ∑_***x***_ *f* (*x*)*P* (*x*),

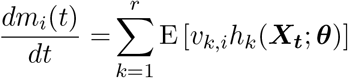

Finally, by linearity of expectation,

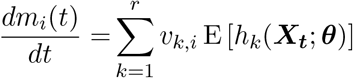

A similar approach can be extended to define a system of ODEs for the time evolution for second order moments of ***X***(*t*),

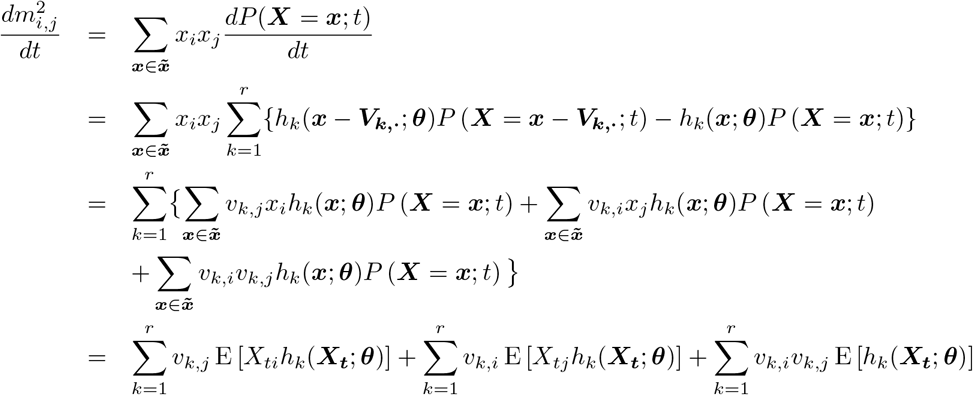

### Supplement B: Example with *N=3* cell types

(http://www.e-publications.org/ims/support/dowload/imsart-ims.zip). In this section the most relevant elements defined in section 3 and section 4 are derived, to allow parameters inference for an illustrative hypothetical *N* = 3 stochastic cell differentiation model. We define the parameters governing stochastic cell differentiation process as

- Individual cell duplication rates vector

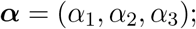
- Individual cell death rates vector:

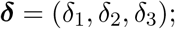
- Individual cell differentiation rates:

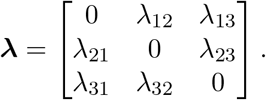

According to the ordering rule described in section 3, the *r* = 12 distinct cellular events are associated with a vector of events rates, ***h***(***X, θ***),

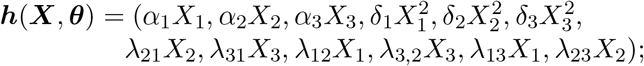

and a net effect matrix ***V***,

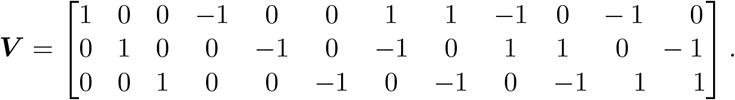

Within the local linear approximation framework described in section Supplement E, the diagonal matrix *D*(***X***) corresponds to

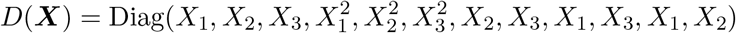

The ODEs systems for time evolutions of process first-order moments is given by

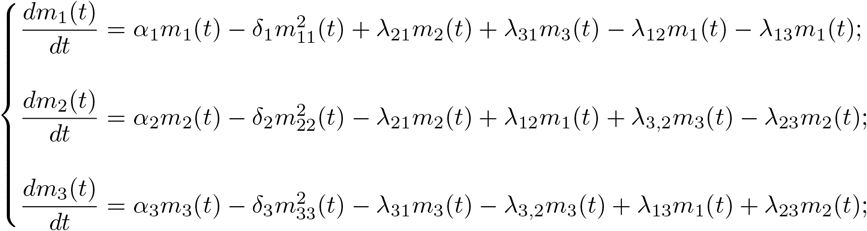

and for second-order moments

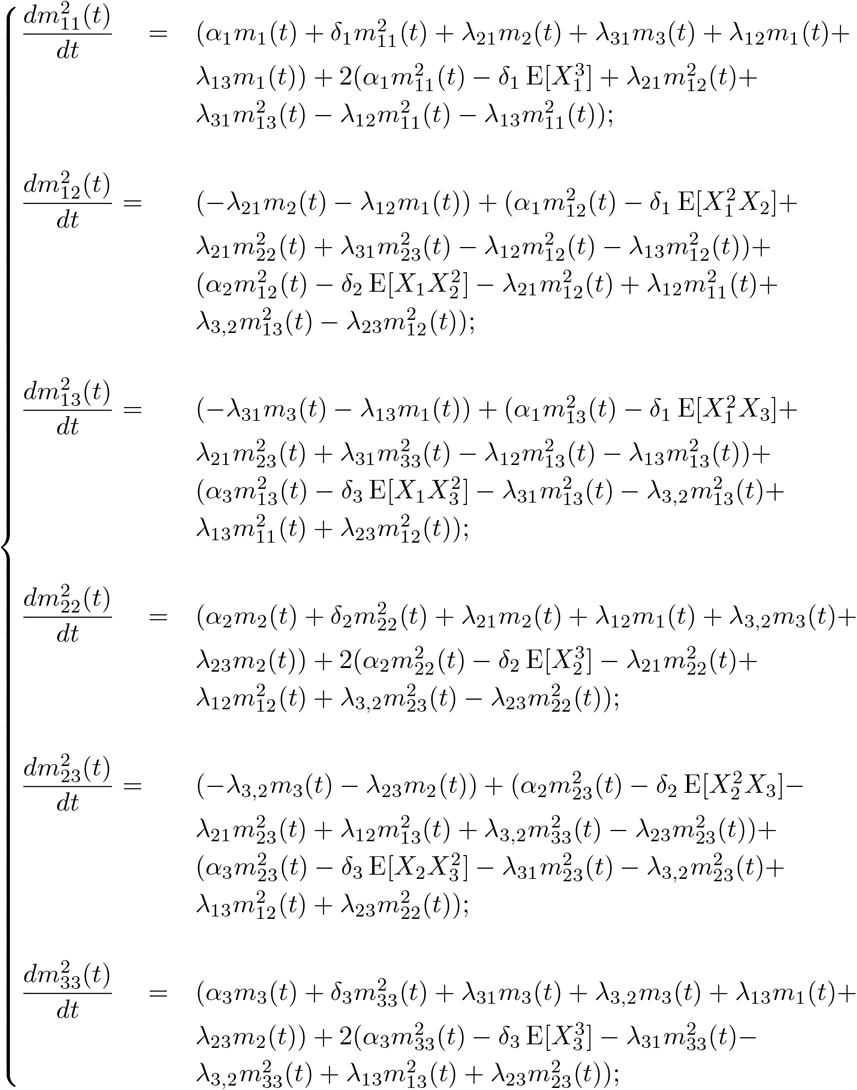

To remove the dependence of second-order moments on higher-order moments, is possible to apply the moment closure schema introduced in section 3.1 and formulated in (3.4).

### Supplement C: Reconstructing cell differentiation network

(http://www.e-publications.org/ims/support/dowload/imsart-ims.zip). In order to investigate the structure of the differentiation tree, differentiation parameters ***λ*** are tested by means of the following asymptotic approximation derived from the generalized method of moments theory (**?**),

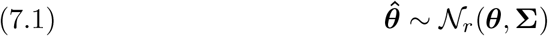

where 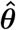is the final vector estimates returned by Algorithm 1 and the asymptotic covariance matrix **Σ** is a *r* × *r* matrix, estimated by means of

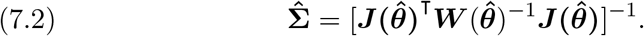

These distributional consideration are used to define Wald-type tests for the differentiation parameters,

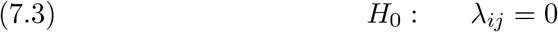

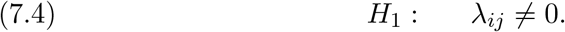

In general, we reject *H*_0_ and conclude that cell type *i* can differentiate into cell type *j*, if 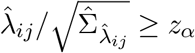. To take into account the positivity constraint, we consider a truncated normal distribution under *H*_0_ as asymptotic distribution, with mean zero and variance equal to the corresponding diagonal element of 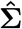and domain restricted to [0, +∞).

### Supplement D: Simulation study with 5 cell types

(http://www.e-publications.org/ims/support/dowload/imsart-ims.zip). In this supplement, we describe the parameter setting used in the simulation study of section 5.1 and shown in Figure 2a. We consider a cell differentiation network with 5 cell types, and therefore 5 cell duplication parameters ***α***, 5 cell death parameters ***δ***, as well as 5 cell differentiation parameters ***λ***:

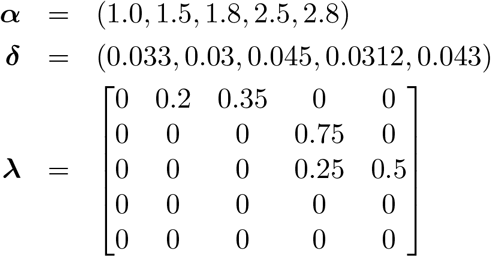

The Gillespie algorithm is implemented in **C++** (Stroustrup, 1997) with the support of **Eigen** library (Guennebaud et al., 2010). Our inferential procedure, described in Algorithm 1, is implemented in **R** (R Core Team, 2015) by means of custom scripts requiring Matrix packages for efficient dense and sparse matrices manipulations Bates and Maechler (2015) and integrated with **C++** scripts calling **ODEint** (Ahnert and Mulansky, 2011) routines that are available in the **Boost** library (Nakariakov, 2013). The quadratic programming problem is solved by means of **IBM ILOG CPLEX Optimizer**, freely available under IBM Academic Initiative program (IBM, 2010). All code used in this manuscript can be found in the online Supplement, and the latest version of the code is available at github.com/dp3ll1n/SLCDP_v1.0.

### Supplement E: Local linear approximation

(http://www.e-publications.org/ims/support/dowload/imsart-ims.zip). In this supplement, we describe a linear approximation of (4.1), which provides quick estimates for the parameters ***θ***. This linear estimate is used in this paper in two different situations. First and foremost, it provides reasonable initial values for the exact non-linear algorithm described in Section 4.1. Secondly, it serves as a comparison in the evaluation of the proposed inference procedure for different sampling intervals. The linear approximation consists of calculating a computationally efficient, albeit approximate, solution for *m*_*i*_(*s*) and 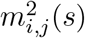 in (**??**) by Euler’s method,

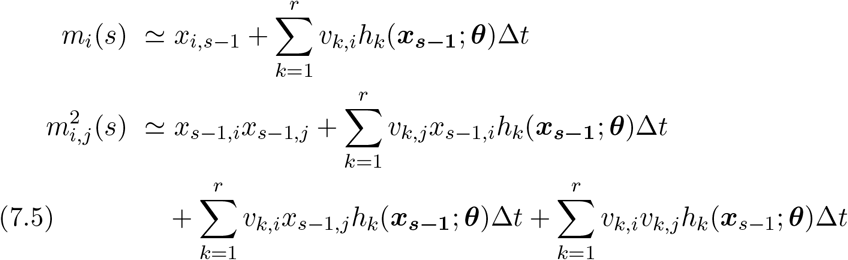

Since (7.5) is linear in ***θ***, the regression model in (4.1) can be conviently reformulated as

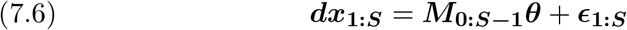

where ***dx***_**1:*S***_ = ***x***_**1:*S***_ − ***x***_**0:*S*−1**_ is column vector with observed cells counts differences between consecutive time points, ***M***_**0:*S*−1**_***θ*** = ***V*** *D*(***x***_**0:*S*−1**_)Δ*t****θ*** is a compact matrix equivalent of (7.5) with *D*(***x***_*s*_) an *r* × *r* diagonal matrix with the appropriate polynomial of ***x***_***s***_ and Var(***E***_*s*_) component is estimated using **Ω**_**0:*S*−1**_ = ***V*** *D*(***x***_**0:*S*−1**_)Δ*t* Diag (***θ***) ***V***. Analogously to (4.2) the local linear estimate 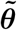 are derived by means of an iterative procedure, in which the following constraint least squares problem is solved,

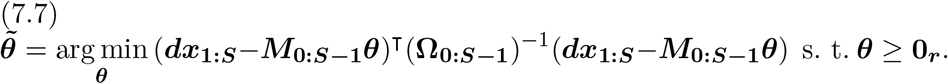

The first estimate 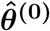, used also as a starting point for the non-linear procedure, is calculated assuming homoscedastic and uncorrelated errors **Ω**_***s***_ = *I*_*N*_ .

### Supplement F: Human hematopoiesis parameter estimates

(http://www.e-publications.org/ims/support/dowload/imsart-ims.zip). In the main paper, we compare a large number of models. For the selected model, we provide here the parameter estimates.

**Table 1.** Parameter estimates for hematopoiesis in human, in-vivo, based on gene therapy clinical trial data, assuming an underlying stochastic cell differentiation process.

| Parameter | Estimate | Parameter | Estimate |
| --- | --- | --- | --- |
| $\alpha_{CD34\_HSC}$ | 8.006e+00 | $\lambda_{CD34\_HSC \rightarrow CD3\_BM}$ | 0.721 |
| $\alpha_{CD3\_BM}$ | 7.930e-01 | $\lambda_{CD34\_HSC \rightarrow CD14\_BM}$ | 0.867 |
| $\alpha_{CD14\_BM}$ | 9.187e-11 | $\lambda_{CD34\_HSC \rightarrow CD15\_BM}$ | 0.591 |
| $\alpha_{CD15\_BM}$ | 3.900e-10 | $\lambda_{CD34\_HSC \rightarrow CD19\_BM}$ | 1.453 |
| $\alpha_{CD19\_BM}$ | 1.251e-01 | $\lambda_{CD34\_HSC \rightarrow CD56\_BM}$ | 0.146 |
| $\alpha_{CD56\_BM}$ | 2.852e-10 | $\lambda_{CD34\_HSC \rightarrow CD61\_BM}$ | 0.335 |
| $\alpha_{CD61\_BM}$ | 5.737e-01 | $\lambda_{CD34\_HSC \rightarrow GLYCO\_BM}$ | 0.713 |
| $\alpha_{GLYCO\_BM}$ | 1.643e-10 | $\lambda_{CD3\_BM \rightarrow CD3\_PB}$ | 0.386 |
| $\alpha_{CD3\_PB}$ | 6.690e-11 | $\lambda_{CD3\_BM \rightarrow CD4\_PB}$ | 0.180 |
| $\alpha_{CD4\_PB}$ | 9.959e-11 | $\lambda_{CD3\_BM \rightarrow CD8\_PB}$ | 0.276 |
| $\alpha_{CD8\_PB}$ | 4.176e-11 | $\lambda_{CD3\_BM \rightarrow CD56\_PB}$ | 0.151 |
| $\alpha_{CD14\_PB}$ | 1.006e-10 | $\lambda_{CD14\_BM \rightarrow CD56\_PB}$ | 0.384 |
| $\alpha_{CD15\_PB}$ | 7.344e-11 | $\lambda_{CD15\_BM \rightarrow CD14\_PB}$ | 0.207 |
| $\alpha_{CD19\_PB}$ | 3.763e-11 | $\lambda_{CD15\_BM \rightarrow CD15\_PB}$ | 0.223 |
| $\alpha_{CD56\_PB}$ | 1.770e-10 | $\lambda_{CD19\_BM \rightarrow CD4\_PB}$ | 0.149 |
| $\delta_{CD34\_HSC}$ | 2.393e-02 | $\lambda_{CD19\_BM \rightarrow CD19\_PB}$ | 0.372 |
| $\delta_{CD3\_BM}$ | 1.735e-04 | $\lambda_{CD19\_BM \rightarrow CD56\_PB}$ | 0.054 |
| $\delta_{CD14\_BM}$ | 2.308e-04 | $\lambda_{CD56\_BM \rightarrow CD56\_PB}$ | 0.153 |
| $\delta_{CD15\_BM}$ | 2.510e-04 | $\lambda_{CD61\_BM \rightarrow CD15\_PB}$ | 0.281 |
| $\delta_{CD19\_BM}$ | 2.322e-03 | $\lambda_{CD61\_BM \rightarrow CD56\_PB}$ | 0.085 |
| $\delta_{CD56\_BM}$ | 6.831e-04 | $\lambda_{GLYCO\_BM \rightarrow CD14\_PB}$ | 0.153 |
| $\delta_{CD61\_BM}$ | 4.734e-03 | | |
| $\delta_{GLYCO\_BM}$ | 1.333e-03 | | |
| $\delta_{CD3\_PB}$ | 1.417e-04 | | |
| $\delta_{CD4\_PB}$ | 3.366e-04 | | |
| $\delta_{CD8\_PB}$ | 2.601e-05 | | |
| $\delta_{CD14\_PB}$ | 1.512e-03 | | |
| $\delta_{CD15\_PB}$ | 5.115e-04 | | |
| $\delta_{CD19\_PB}$ | 4.390e-04 | | |
| $\delta_{CD56\_PB}$ | 2.630e-04 | | |

### Supplement G: Parameter estimates sensitivity to random initialization

(http://www.e-publications.org/ims/support/dowload/imsart-ims.zip). Here we show the sensitivity of the estimates to random initializations.

